# Effects of auditory sleep modulation approaches on brain oscillatory and cardiovascular dynamics

**DOI:** 10.1101/2022.02.16.480303

**Authors:** Stephanie Huwiler, Manuel Carro Dominguez, Silja Huwyler, Luca Kiener, Fabia Stich, Rossella Sala, Florent Aziri, Anna Trippel, Christian Schmied, Reto Huber, Nicole Wenderoth, Caroline Lustenberger

**Affiliations:** Neural Control of Movement Lab, Institute of Human Movement Sciences and Sport, Department of Health Sciences and Technology, ETH Zurich, 8092 Zurich, Switzerland; Department of Cardiology, University Heart Center Zurich, University of Zurich, Zurich, Switzerland; Center of Competence Sleep & Health Zurich, University of Zurich, Zurich, Switzerland; Neuroscience Center Zurich (ZNZ), University of Zurich, ETH Zurich, Zurich, Switzerland; Child Development Centre, University Children’s Hospital, University of Zurich, Zurich, Switzerland; Department of Child and Adolescent Psychiatry and Psychotherapy, Psychiatric Hospital Zurich, University of Zurich, Zurich, Switzerland; Future Health Technologies, Singapore-ETH Center, Campus for Research Excellence and Technological Enterprise (CREATE), Singapore

## Abstract

Sleep modulation techniques to elucidate the functional role of sleep brain oscillations in brain and body functions have gained large interest. Slow waves, the hallmark feature of deep non-rapid eye movement sleep, do potentially drive restorative effects on brain and cardiovascular functions. Auditory stimulation to modulate slow waves is a promising tool, however, directly comparing different auditory stimulation approaches within a night and analyzing induced dynamic brain and cardiovascular effects are yet missing. Here, we tested various auditory stimulation approaches in a windowed, 10 s ON (stimulations) followed by 10 s OFF (no stimulations), within-night stimulation design and compared them to a SHAM control condition. We report the results of three studies and a total of 51 included stimulation nights. We found a large and global increase in slow wave activity (SWA) in the stimulation window compared to SHAM. Furthermore, slow wave dynamics were most pronouncedly increased at the start of the stimulation and declined across the stimulation window. Beyond the changes in brain oscillations, we observed, for some conditions, a significant increase in the mean interval between two heartbeats within a stimulation window, indicating a slowing of the heart rate, and increased heart rate variability derived parasympathetic activity. Those cardiovascular changes were positively correlated with the change in SWA and thus, our findings provide mechanistic insight into the potential of auditory slow wave enhancement to modulate cardiovascular restorative conditions during sleep. However, future studies need to investigate whether the potentially increased restorative capacity through slow wave enhancements translates into a more rested cardiovascular system on the subsequent day.

## Introduction

Sleep represents a powerful system for promoting brain and body health. It is suggested to play a role in a plethora of functions such as cleaning away toxic by-products (Eide et al., 2021; Eugene & Masiak, 2015; Underwood, 2013), synaptic homeostasis(Tononi & Cirelli, 2006), memory consolidation (Leminen et al., 2017; Hong-Viet V. Ngo, Martinetz, et al., 2013; Ong et al., 2016, 2018; Papalambros et al., 2017; Santostasi et al., 2015; Simor et al., 2018), metabolic (D. J. Dijk, 2008), and cardiovascular functions (Knutson et al., 2007; Mullington et al., 2009; Palagini et al., 2013; Xi et al., 2014), and body core tissue turnover (Stich et al., 2021). Particularly non-rapid eye movement (NREM) sleep with its large-amplitude, low-frequency slow waves has been proposed to guide those beneficial effects (e.g., reviewed in (Bellesi et al., 2014)). Periods of neuronal activity are reflected in the slow wave up-phase and periods of neuronal silence are reflected by the down-phase of slow waves (Steriade et al., 1993), thereby coordinating the temporal interplay between thalamocortical sleep spindles and hippocampal sharp wave ripples, which has for instance been shown to support long-term memory retention (Diekelmann & Born, 2010; Helfrich et al., 2018). Nevertheless, whether slow waves are an indispensable driver for the maintenance of a healthy brain and body remains still largely unexplored.

To elucidate the functional role of slow waves for brain and body functions, modulation of these oscillations is needed. Over the last few years, especially auditory stimulation has emerged as a promising, non-invasive, and feasible approach to selectively modulate slow waves during deep sleep (Besedovsky et al., 2017; Grimaldi et al., 2019; Hong-Viet V. Ngo, Martinetz, et al., 2013). However, various stimulation protocols exist, leading to inconsistent findings on behavioral outcomes (e.g., reviewed in (Wunderlin et al., 2021)) and comparisons of those approaches on efficacy to selectively enhance or decrease slow waves are missing. Ngo and colleagues (Hong-Viet V. Ngo, Martinetz, et al., 2013) were the first to report that targeting the ascending up-phase of ongoing slow waves seems to be important to elicit improvements in overnight memory consolidation. Down-phase stimulation on the other hand was shown to rather interfere with slow waves and the consolidation of declarative and motor memory (Fattinger et al., 2017; Hong-Viet V. Ngo, Martinetz, et al., 2013). However, in addition to selecting an appropriate target phase of the auditory stimuli, the number of stimulations in a sequence is variable, as e.g., a two tones stimulation protocol with a subsequent stimulation break afterwards (Besedovsky et al., 2017; Hong-Viet V. Ngo, Martinetz, et al., 2013), or a windowed approach where auditory stimulation was only presented during an ON window of a predefined length (Grimaldi et al., 2019; Papalambros et al., 2017; Santostasi et al., 2015). Besides the mentioned procedures that rely all to a certain degree on the phase and/or presence of slow waves (closed-loop stimulation), completely open-loop auditory stimulation has been shown to enhance slow waves as well (Simor et al., 2018; Weigenand et al., 2016). An additional parameter to consider is the volume of stimulation and whether the stimuli are played through headphones or with loudspeakers. Additionally, some studies used fixed volumes between 50 - 60 dB (Besedovsky et al., 2017; Hong-Viet V. Ngo, Claussen, et al., 2013; Hong-Viet V. Ngo, Martinetz, et al., 2013), or individual and/or adaptive volumes ranging between 30 - 60 dB (Grimaldi et al., 2019; Leminen et al., 2017; Simor et al., 2018).

Although many stimulation approaches have been applied, auditory stimulation is still in its infancy. Thus, the full potential of auditory stimulation has not been exploited and more profound understanding of its effects is needed for this purpose. Furthermore, it is currently unknown whether auditory stimulation efficacy remains stable across a sleep cycle and whether the stimulation efficacy is even maintained across several seconds of stimulation. To advance the understanding of auditory slow wave modulation we propose here a novel approach to compare different auditory stimulation conditions within a single sleep period using a windowed 10 s stimulation ON (auditory stimulation played) followed by 10 s OFF (no auditory stimulation played) approach. This within-night design eliminated any between-night variability, which is observed as pronounced fluctuations of for instance sleep architecture or depth of NREM sleep across multiple nights within the same subject. Therefore, our set-up allows for controlled comparisons between conditions. Because the ON window only starts if a set of stimulation prerequisites such as stable NREM sleep are met and a within-night SHAM control can be implemented, our design allows for a direct comparison of stimulation effects using a larger parameter space. Furthermore, this windowed approach entails many advantages compared to continuous approaches as underlying temporal dynamical brain and body responses can be analyzed. Thus, averaging across a longer time period can be avoided. Furthermore, such a windowed approach had recently been shown to be more efficient (Lustenberger et al., under review) in enhancing SWA compared to a continuous approach, however, this has been shown between nights only.

Beyond the elicited responses of the auditory stimulation on brain activity, cardiovascular parameters such as five minutes heart rate variability (HRV) (Grimaldi et al., 2019) were shown to be altered upon slow wave enhancement. During sleep, the heart is mainly under the control of the autonomic nervous system (ANS) dominated by a cyclic pattern with parasympathetic predominance during NREM sleep (Simon et al., 1998) and sympathetic predominance during rapid eye-movement (REM) sleep (Somers et al., 1993). The strong positive correlation between slow wave activity (SWA, power spectral density in the slow wave frequency band) and parasympathetic activity (Jurysta et al., 2003) indicates a link between slow waves and the resting function of the parasympathetic nervous system. However, whether slow waves play a functional role in the restorative parasympathetically dominated control of the cardiovascular system remains to be elucidated. Human ANS activity, and especially parasympathetic activity, can be indirectly derived by HRV parameters (Schlosser, 1996) that are calculated based on the intervals between successive normal heart beats (RR intervals). However, the ECG signal from which the HRV parameters are derived is continuous. Thus, temporarily more precise measures such as instantaneous heart rate (IHR, inverse of RR intervals) may suit better to reflect immediate cardiovascular dynamics as an outcome of altered cardiac autonomic regulation upon auditory slow wave modulation.

Here, we (i) compared widely used auditory stimulation approaches on their potential to modulate slow waves and (ii) investigated whether auditory stimulation modulates not only slow waves but also cardiovascular dynamics. Additionally, we explored the effects of auditory stimulations on sleep stage shifts and arousals and tested whether the timing of the stimulations within a night influences the effects on slow wave modulation by splitting the night into the first four hours of stimulation and the remaining hours of stimulation. Altogether, we aimed to provide a framework about efficient stimulation conditions for future studies investigating the functional role of slow waves for the brain and body.

## Methods

### Participants

Overall, 63 healthy male participants were enrolled in three studies (Study1: n = 23, age = 40.25 ± 13.69 years, Study2: n = 9, age = 50.36 ± 6.74 years, Study3: n = 31, age = 45.48 ± 9.78 years) that served as preparation studies to optimize the auditory stimulation settings of a main trial (registered at ClinicalTrials.gov (NCT04166916)). All participants were non-smokers, reported a regular sleep-wake rhythm, a body-mass index between 17 - 30, and none of the participants had a presence of psychiatric/neurological diseases, presence or suspicion of sleep disorders, or presence of clinically significant concomitant diseases. Participants taking on-label sleep medication or medication affecting the cardiovascular system (e.g., beta-blockers) were further excluded. We recruited the participants from the community using advertisements on different platforms and were supported by the ETH Alumni Association. The study was approved by the cantonal Ethics Committee Zurich. All participants provided written informed consent before participation and received monetary compensation for their participation. For this analysis, we excluded seven participants of Study3 because of either very poor visually detected sleep quality in the nightly spectrogram (n = 4: e.g., almost complete night covered with alpha activity), severe sleep apnea (n = 2), and high blood pressure (n = 1) meaning direct exclusion for the main trial and thus, no sleep scoring was performed. Because Study3 was a proof-of-principle about a dose-dependency effect of volume of stimulation and to directly compare those volumes (see section auditory stimulation conditions), we chose a subset of participants from this study in order to maximize the effect while using identical volumes for all participants. Thus, we decided to exclude five participants that did not show a Hilbert Amplitude response to any of the sound volumes to isolate the effect in responders. Why those participants did not show a response to the tones remains to be explored and personalized sound settings may help to overcome this issue. Therefore, the included sample size of Study3 is n = 19 (age: 45.29 ± 11.00 years). Together with the participants of Study1 (n = 23) and Study2 (n = 9), we included in total 51 nights in this analysis.

### Experimental procedure

All participants underwent a screening period of a first phone screening where we verified the eligibility of participants. Thereafter, participants were invited to our sleep laboratory. All participants were asked to adhere to a regular sleep rhythm within two nights before the screening night, abstain from caffeine after 15:00, naps, alcohol, and sports on the day of the screening night. Compliance was controlled by sleep logs and questionnaires. After arrival at our sleep lab, participants underwent a basic physiological assessment by measuring height, weight, waist circumference, and hip circumference. Thereafter, participants answered several questionnaires about demographics, health status, handedness, chronotype, sleep habits and sleep quality, and daytime sleepiness. However, we do not report the outcomes of the questionnaires as it is beyond the scope of this paper. We attached the high-density EEG system (Geodesic Sensor Net, Magstim EGI, Eugene, USA) and the ECG electrodes with a modified lead II Einthoven configuration using gold recording electrodes (Genuine Grass electrodes, Natus Medical Inc., Pleasanton, US). Afterwards, we performed a simple audiometry to ensure participants can perceive a sound volume of 45 dB. All participants were able to hear 40 dB tones in both ears, except one only correctly perceived 40 dB in one ear. However, this participant was part of Study1 where only 45 dB sounds were played during the night. All impedances were kept below 40 kΩ. Finally, we attached a breathing belt and a canula for detecting sleep apnea (BrainProducts GmbH, Gilching, Germany and SOMNOmedics GmbH, Randersacker, Germany). Participants with clear sleep apnea were excluded from subsequent analyses. Participants were allowed to sleep for 8 hours according to their normal bedtime while we recorded polysomnography (BrainProducts GmbH, Gilching, Germany) and ECG at a sampling rate of 500 Hz. Furthermore, we recorded continuous blood pressure (Finapres Novascope, Finapres Medical Systems BV, Enschede, the Netherlands and SOMNOmedics GmbH, Randersacker, Germany), however, blood pressure data is not reported here.

### Auditory stimulation protocol

To reliably compare auditory stimulation approaches we designed a novel stimulation protocol based on the algorithm of a previously established mobile stimulation device (Ferster et al., 2019) that was adapted for a high-density in-lab solution using OpenViBE (Renard et al., 2010). Briefly, a single-channel EEG input (Fpz-A2) was used and processed to trigger the auditory stimulation during stable NREM sleep. The complete procedure on the stimulation logic is described in detail here (Lustenberger et al., under review). Briefly, the EEG signal was notch filtered at 50 Hz, then the EEG spectral power was calculated in the three frequency bands: low delta (2-4 Hz), high delta (3-5 Hz), and high beta (20-30 Hz) to determine whether the participant is in NREM sleep or non-NREM sleep (wake or REM). Furthermore, the first 10 minutes of uninterrupted NREM sleep served to calculate a beta power threshold for each night to account for a shift to wake, light sleep, or artifacts. Thus, if after the first 10 minutes beta power exceeded this threshold, stimulation was halted. To improve accuracy of stimulations during NREM sleep, we implemented a correlation index for two electrooculography deviations that interrupted sleep detection if a high anti-correlation was measured. Additionally, we implemented a movement detection based on the absolute amplitude of the EEG signal to delay stimulations for 10 seconds after a movement has been detected. Lastly, we implemented a slow wave sleep detection algorithm based on the power in the delta frequency range to only stimulate when delta power exceeded a predefined threshold. To deliver stimulations at a specific phase of ongoing slow waves, a first-order phase-locked loop (PLL) was implemented according to Ferster et al. (in preparation), that was initially adapted from Santostasi et al. (Santostasi et al., 2015). We set the target phase of the PLL to 50° for up-phase stimulations and 230° for down-phase stimulations. Auditory stimulations were presented through Etymotic insert earphones (Etymotic Research Inc., ER 3C).

The stimulation conditions were presented in a pseudo-randomized order in a windowed ONOFF approach during detected periods of stable NREM sleep by the algorithm, as illustrated in Figure 1. Every stimulation ON window started after the target phase of the PLL has been reached (up-phase for all conditions except down-phase stimulations). Afterwards during the 10 s ON window auditory stimuli were presented. We chose this stimulation duration because any change in cardiac autonomic modulation may need more than five seconds to occur depending on parasympathetic or sympathetic modulation (Nunan et al., 2010). After the ON window, the 10 s OFF window started where only triggers were saved but no auditory stimulation was applied. Note that for the SHAM control condition, auditory stimulation was applied in neither ON nor OFF windows, however, triggers were saved for both. All conditions were randomly shuffled and presented in that sequence, whenever stimulation criteria were met. Only after every condition within a sequence was presented, the order of the conditions was re-shuffled again for the complete sleep period of 8 hours. Note that the ONOFF window together will be referred to as stimulation window.

**Figure 1:**
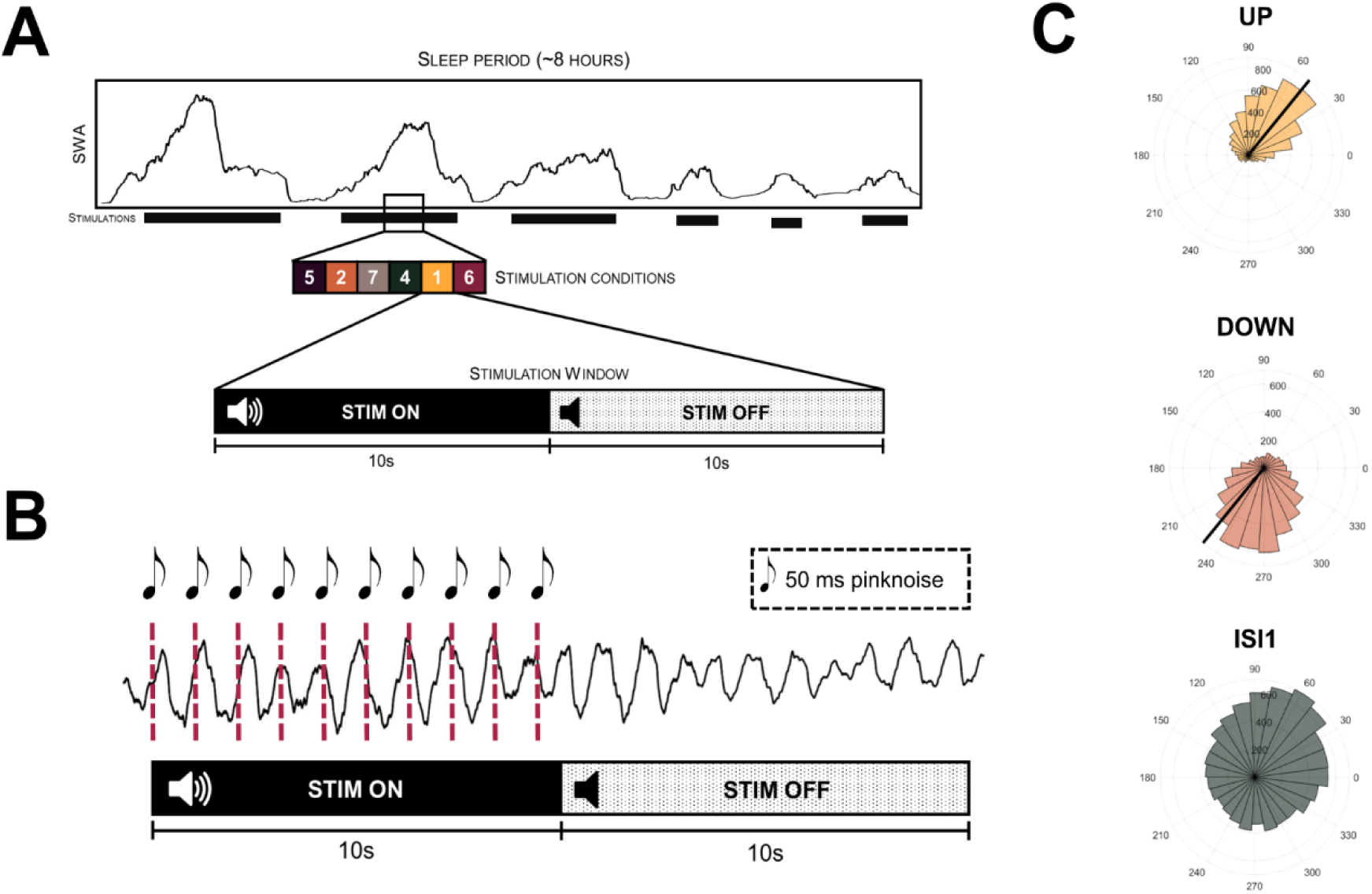
Experimental design. A: Stimulation design of within night presentation of different auditory stimulation conditions. Whenever the automatic sleep detection algorithm during the complete sleep period of 8 hours detected stable non-rapid eye movement sleep and the phase-locked loop reached the threshold target phase, a stimulation window was triggered. A single stimulation window consists of 10 s stimulation ON, where auditory stimulations were applied according to the presented condition, directly followed by a stimulation OFF window, where no auditory stimulation was presented. All conditions were presented in pseudo-randomized order, meaning that they got shuffled in the beginning and only after all different conditions have been presented, the order got reshuffled again. Note that during the SHAM condition, no auditory stimulations were presented in neither the ON nor the OFF window. B: Exemplary stimulation (rhythmic 1 Hz stimulation, ISI1) for a single stimulation window. C: Circular distribution of the real target phase of the auditory stimulation for the conditions UP (ascending phase of all detected slow waves was targeted), DOWN (descending phase of all detected slow waves was targeted), and ISI1 stimulation for n = 23 participants of Study1.

### Auditory stimulation conditions

We compared various auditory stimulation approaches that are all described in Table A1 as an overview. Here, we report the results of the most comparable approaches that fall in the following categories: Phase-specific, rhythmic, and sound modulated.

Previous research has shown that down-phase auditory stimulation suppresses slow waves (Fattinger et al., 2017; Hong-Viet V. Ngo, Martinetz, et al., 2013), whereas up-phase stimulation has repeatedly been shown to increase slow wave activity (Besedovsky et al., 2017; Hong-Viet V. Ngo, Martinetz, et al., 2013; Papalambros et al., 2019). We employed a condition to target the ascending phase of slow waves (UP) and a condition to target the descending phase (DOWN) of slow waves and only detected slow waves by the algorithm were stimulated during the stimulation ON window. All auditory stimuli consisted of a 50 ms burst of pink noise.

To elaborate on the stimulation protocol used by Ngo and colleagues (Hong-Viet V. Ngo, Martinetz, et al., 2013) where they targeted the first ascending phase of a slow wave and presented the second tone one cycle of a slow wave later, we extended this protocol to a continuous rhythmic stimulation to target the first ascending-phase of a slow wave and continuing in a 1 Hz rhythm throughout the stimulation ON window (ISI1). Lastly, we tested two more sophisticated conditions: 1 Hz amplitude modulated pink noise (ENVELOPE) and BINAURAL BEATS with a difference of 1 Hz between the two ears and carrier frequency of 400 Hz. Both methods should elicit entrainment in the target frequency range (Chaieb et al., 2015; Lustenberger et al., 2018) of 1 Hz and therefore, increase power spectral density in the slow wave frequencies.

During Study1, we kept the sound volume constant at 45 dB, afterwards we specifically investigated the effects of various sound volumes and sound volume modulation on changes in EEG and ECG. In Study2, we tested the effects of slowly increasing and decreasing sound volume during ISI1 stimulation. More precisely, the sound volumes of the ISI1_Mod_ condition (reported in dB) were as follows: 40, 40, 42.5, 42.5, 45, 45, 45, 45, 42.5, 42.5. To further investigate the volume effects, we conducted the final Study3 employing the ISI1 condition with different sound volumes. (1) ISI1_High_: constant 45 dB and (2) ISI1_Low_: constant 42.5 dB.

In addition to the above reported conditions, we tested four more conditions within the three studies. We do not report the results of those conditions here because they either provide no additional information for our main research question or are highly explorative paradigms (e.g., not used in previous studies).

### EEG analysis and sleep scoring

EEG analyses were performed using the EEGLAB toolbox (Delorme & Makeig, 2004) in MATLAB (R2019a, MathWorks Inc., Natick, MA), and pre-processing consisted of resampling to 200 Hz and using the PREP pipeline (Bigdely-Shamlo et al., 2015) to remove line-noise, robust average referencing, and bad channel interpolation. Afterwards, data was band-passed filtered between 0.5 and 40 Hz using FIR filters. Artifacts were rejected based on a semiautomatic artifact removal procedure that was described previously (Lustenberger et al., 2015). Additionally, we applied a previously published automatic arousal detection to the Fz, Pz, and Oz channels (Alvarez-Estevez & Fernández-Varela, 2019; Fernández-Varela et al., 2017). This algorithm uses one EMG derivation and one EEG derivation to detect alpha and beta arousals (Alvarez-Estevez & Fernández-Varela, 2019; Fernández-Varela et al., 2017). Sleep was scored visually for sleep stages in 20 seconds epoch according to standard criteria (Iber et al., 2007) by one expert scorer. Only stimulation windows in artefact and arousal-free N2 or N3 windows were included in further analyses. A fast Fourier transform was applied using the MATLAB *pwelch* function with a Hanning window of 4 seconds with 50 % overlap to calculate the power spectral densities (PSD) in the frequency bands of interest. To compare the PSD distribution for the stimulation windows, we either normalized the complete power spectra by its cumulative power up to 30 Hz (e.g., applied in (Hong-Viet V. Ngo, Martinetz, et al., 2013; Ong et al., 2016)) and for comparison the PSD without normalization because of the large increase in the SWA bands covering all changes in other frequency bands as well. Thereafter, the relative change was calculated by applying the following formula: 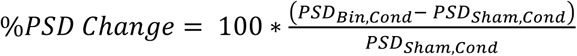. To analyze the phase of the stimulation and the temporal EEG response we first band-pass filtered the EEG data using a Hamming windowed sinc FIR filter with the EEGLAB function *pop_eegfiltnew* in the respective frequency bands and thereafter, Hilbert transformed the data to extract the Hilbert phase and amplitude. We will refer to the Hilbert amplitude as slow wave dynamics.

To elucidate whether power changes in the spindle frequency band are driven by sleep spindles or are rather accompanied by arousals because of the overlapping frequency bands, we detected all sleep spindles across all electrodes using an established algorithm (Ferrarelli et al., 2007, 2010). Briefly, we first down-sampled the signal to 200 Hz, and band-pass filtered the signal between 10 to 16 Hz using a Chebychev filter and the MATLAB function *filtfilt*. Afterwards, sleep spindles were detected with a threshold of two times the mean to five times the mean of the filtered signal. The spindle frequency and timepoint were derived for all detected spindles. Thereafter, we assigned for bins of 2 seconds of all stim windows for each participant a value of either 1 if a detected spindle started or 0 if no spindle start was detected in the respective bins. Finally, we calculated a probability distribution by dividing the sum of all detected events by the total amount of valid stimulation windows for each bin.

### ECG Analysis

ECG R peaks were automatically detected and visually corrected if necessary using PhysioZoo (Behar et al., 2018). Segments of non-detectable R peaks or poor data quality were marked and excluded for further analyses. The generated RR intervals were processed with MATLAB (R2019a, MathWorks Inc.) and only RR intervals with all R peaks occurring completely in a stimulation window were extracted and the root mean square of successive RR differences (RMSSD) and the standard deviation of normal RR intervals (SDNN) was calculated. Furthermore, IHR at a specific time point was calculated by dividing 60 through the RR interval. Afterwards, continuous IHR was derived by interpolating missing datapoints using the function *fillmissing* with the method *makima*. Because we observed that the interpolation of the last beats in the stimulation window performed poorly, we excluded the last two seconds for the IHR plots and therefore only show the results of the complete ON and eight seconds OFF window. To calculate the relative IHR, we calculated the mean HR of the closest SHAM window if both windows occurred within five minutes, and used the following formula for every timepoint: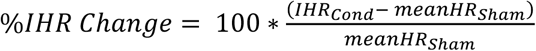. If no SHAM window occurred within five minutes, the window was discarded. Additionally, we extracted the following features for every stimulation window: longest RR interval, shortest RR interval, mean RR interval, and the difference between the longest and shortest RR interval. We will refer to the IHR and HRV measurements as cardiovascular dynamics.

### Statistics

Statistical analyses were conducted in RStudio (R Core Team, 2013) version 1.2.5033 and MATLAB (R2019a). To compare the number of stimulation windows for the first four hours of stimulations to the remaining hours of stimulation, the number of stimulations within a stimulation window for the UP and DOWN condition and their differences for the first four hours of stimulations to the remaining hours of stimulation, we employed paired two-sided t-tests. Further comparisons of two conditions for instance for volume effects were based on paired t-tests as well. To compare the topographical distribution of the EEG power spectral density within specific frequency bands, we ran linear mixed-effects models for each electrode with the fixed factor condition and random factor participant using the MATLAB function *fitmod*. Post-hoc p-values for the fixed effects between the auditory stimulation conditions and the SHAM condition were obtained using the function *fixedEffects* with the Satterwhaite approximation for the degrees of freedom and were corrected for multiple comparisons by false discovery rate (Benjamini & Hochberg, 1995) for each condition and all electrodes. For the Hilbert amplitude analysis, we ran linear mixed-effects models for every timepoint using *fitmod* and factor condition as fixed effects and participant as random effect. Post-hoc p-values for each condition compared to SHAM were obtained using *fixedEffects* with the Satterwhaite approximation for the degrees of freedom. All time course plots show the mean ± standard error of the mean (sem) and were generated in MATLAB using the *boundedline* function (Kearney, n.d.). The HRV and HR values were compared using a linear mixed-effects model with fixed factors Condition and random factor Subject using the software package lme (Bates et al., 2015) in R. If the linear mixed-effects model was significant for the dependent variable (condition), we derived post-hoc p-values using the Satterwhaite method of the package lmerTest (Kuznetsova et al., 2020). Note that we did not expect all conditions to evoke similar effects on the brain and the body and therefore, we also calculated post-hoc p values if the p-value of the model showed trend level (p < 0.1). To correct for multiple comparisons, we used the R package emmeans (Searle et al., 2021), which uses the population marginal means, and the Hochberg method to adjust post-hoc p-values. To directly compare the different volume conditions with each other, we employed two-sided paired t-tests. To further analyze the relationship between the change in SWA with the change in HR(V) features, we employed repeated measures correlations using the R package rmcorr (Bakdash et al., 2016). P-values < 0.05 were considered significant, p values < 0.1 were considered as trend level. Plots were generated using the ggplot2 package of R (Wickham, 2016) or MATLAB.

## Results

We report the results of three studies where we investigated the effects of different auditory stimulation approaches on brain oscillations and cardiovascular dynamics. We separated the different approaches in effects of phase, effects of sound modulation, effects of volume, and effects of time of the night. We first report how these settings affect slow wave dynamics, spectral density, arousals, and spindles, thereafter the effects on cardiovascular dynamics, and finally how slow wave and cardiovascular effects are correlated. Basic sleep architecture parameters for all reported studies are shown in Table A1 in the Appendix.

### Effects of phase targeted auditory stimulation on EEG power spectral density and slow wave dynamics

First, we analyzed the performance of our algorithm to hit a specific phase of a slow wave in comparison with the rhythmic ISI1 protocol. The algorithm showed a high precision in hitting the up-phase or down-phase of all detected slow waves in the STIM ON window as illustrated in Figure 1C. The ISI1 approach led to a distribution of the stimulations across the complete cycle of slow waves. As a next step, we counted the number of stimulations within the stimulation windows. For the UP condition, a mean number of 6.28 ± 0.21 of stimulations and for the DOWN condition, a mean number of 6.33 ± 0.21 stimulations occurred within each window. First, we compared the non-normalized change in power spectral density distribution as illustrated in Figure A1A in the Appendix. We observed particularly strong enhancements in the slow wave frequency bands, however, also increases in other frequency bands were present. Comparing the change in power spectral density distribution that was normalized to the cumulative power from 0.5 - 30 Hz (see Figure A1B in the Appendix), only the slow wave frequency band was increased compared to SHAM, whereas the higher frequencies were reduced. This clearly shows the large power increase in the slow wave frequency bands induced by the auditory stimulations. Because of the strong effects in the slow wave range, we next focused on slow wave dynamics. We analyzed the Hilbert amplitude in the low slow wave frequency band (0.5 - 2 Hz) during the stimulation windows. Figure 2A illustrates the percentage change in slow wave Hilbert amplitude of the Fz electrode to SHAM and shows a clear increase for the UP, DOWN, and ISI1_High_ conditions at the start of the stimulation that is, however, gradually attenuated during the stimulation period. The initial peak represents a mean increase of 81.96 ± 14.99 % (UP), 91.59 ± 13.70 % (DOWN), and 102.05 ± 15.90 % (ISI1_High_) compared to SHAM. As a next step, we investigated whether there was a difference in these maximal Hilbert amplitudes between all non-SHAM conditions. Thus, we ran a linear mixed-effects model of all conditions of Study1 (UP, DOWN, ISI1_High_, ENVELOPE, BINAURAL BEATS) that revealed a significant effect of condition (F_Cond_ (4,88) = 5.79, p_condition_ < 0.001) on the maximal relative Hilbert amplitude compared to SHAM. However, post-hoc comparisons of the conditions UP, DOWN, and ISI1_High_ showed no significant difference between any of these conditions (all p > 0.1).

**Figure 2:**
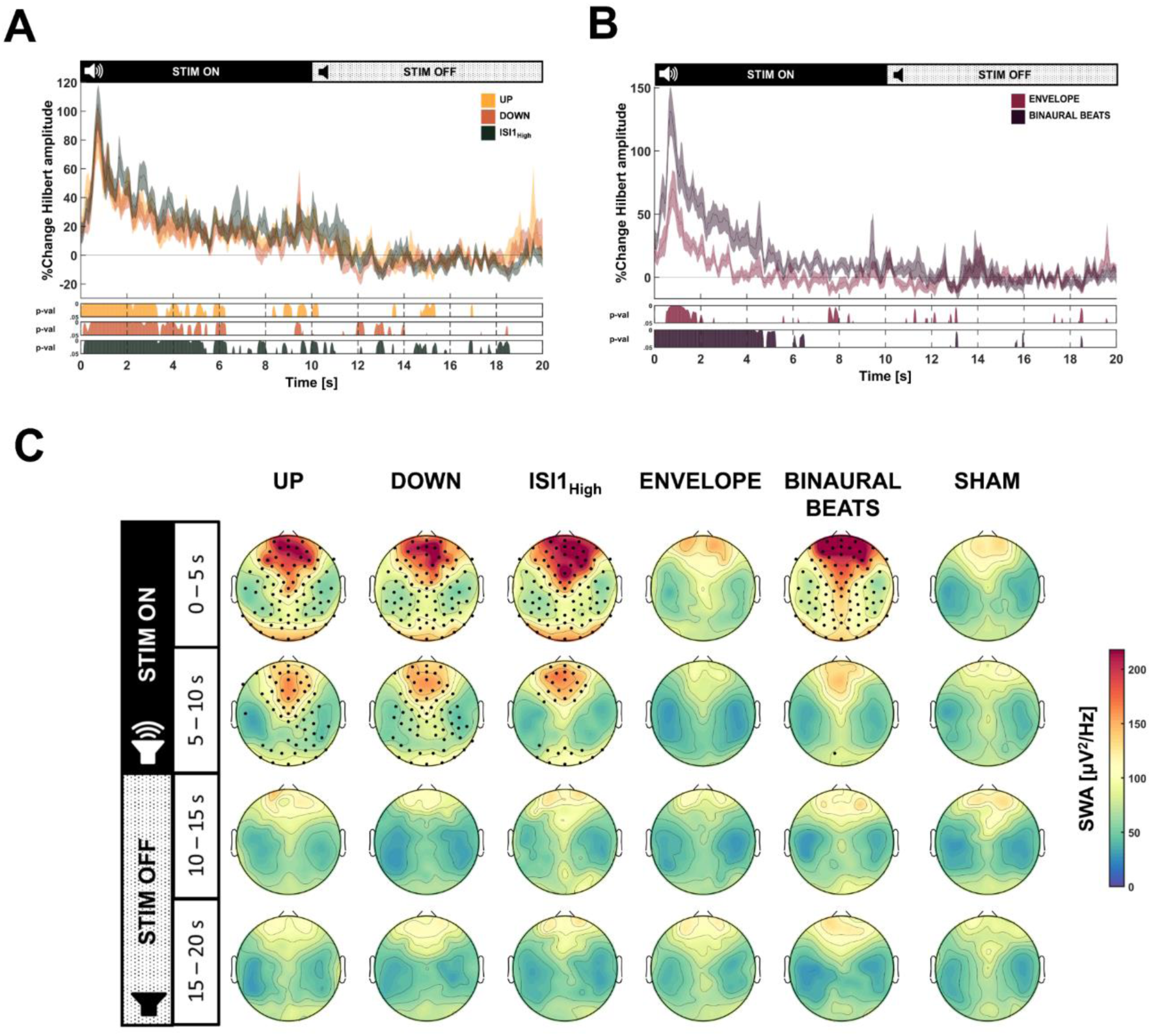
Results of auditory stimulation conditions on slow wave dynamics and slow wave activity. A + B: Percentage change in Hilbert amplitude in the slow wave frequency band (0.5 – 2 Hz) for the conditions UP, DOWN, ISI1_High_ relative to SHAM (A), or ENVELOPE and BINAURAL BEATS relative to SHAM (B) respectively, during the complete stimulation windows. Hilbert amplitude change is presented as mean ± standard error of the mean. The horizontal line at zero represents no change compared to the SHAM condition. Below the plot the resulting p-values of post-hoc comparisons of linear mixed-effects model with condition as independent factor and subject as random factor for every timepoint, is shown for each condition for yellow (UP), orange (DOWN), dark green (ISI1_High_) (A) or pink (ENVELOPE) and dark violet (BINAURAL BEATS) (B) relative to SHAM. Note that effect is most pronounced during beginning of stimulation ON window and diminishes with time. C: Topographical change of slow wave activity (SWA, 0.5 – 2 Hz) for the conditions UP, DOWN, ISI1_High_, ENVELOPE and BINAURAL BEATS compared to SHAM. Because of the difference of the SWA response seen in the Hilbert response, we divided the stimulation ON and OFF window in two 5 s windows each. Black dots indicate significant electrodes (p < 0.05) for the post-hoc p-values resulting from linear mixed-effects effects model models with condition as independent factor and subject as random factor. P-values for each topoplot have been corrected for multiple comparisons by applying the false discovery rate. All plots are shown for n = 23 participants.

The maximal relative increase was 81.96 ± 14.99% (UP), 91.59 ± 13.70% (DOWN), and 102.05 ± 15.90% (ISI1_High_) compared to SHAM, and the respective p-values of the post-hoc comparisons are illustrated below in Figure 2A. Given the non-stationary effect of the auditory stimulation, we next analysed the topographical effect in the low SWA (0.5 - 2 Hz) for every five seconds of the stimulation windows, which is illustrated in Figure 2C. We found a significant increase in SWA across all cortical regions. The change in SWA compared to SHAM is illustrated in Figure A2C in the Appendix. Furthermore, as already expected based on the Hilbert amplitude results, the SWA enhancing effect was most pronounced during the first five seconds of the stimulation ON window. In contrast to previous studies (Fattinger et al., 2017; Hong-Viet V. Ngo, Martinetz, et al., 2013) we did not observe a decrease in SWA upon down-phase stimulation. However, as seen in Figure A2B in the Appendix, there was a significant decrease in high SWA (2.25 - 4.5 Hz) in a central cluster of electrodes in the OFF window of all auditory stimulation conditions (UP, DOWN, ISI1_High_).

### Effects of sound modulation on EEG power spectral density and slow wave dynamics

Different than the phase targeting conditions where only bursts of pink noise were played, we applied two sound modulation conditions based on 10 seconds continuous sounds (either amplitude modulated pink noise (ENVELOPE) or continuous single-frequency sound with a 1 Hz discrepancy between the ears (BINAURAL BEATS)). Figures A1C and A1D show the power spectral density distribution change compared to the SHAM condition. We observed a strong increase in the slow wave frequency bands of the non-normalized distribution for the BINAURAL BEATS, whereas the increase in the low frequencies was not as pronounced for the ENVELOPE condition. Furthermore, within the BINAURAL BEATS, the percentage power change of the lower alpha bands was comparable to the percentage power increase of the slow wave range. Similar to the phase-targeting conditions, we also investigated the slow wave dynamics (Hilbert amplitude of Fz, 0.5 - 2 Hz) which is shown in Figure 2B as the percentage change compared to SHAM. The initial peak represents a mean increase of 131. 30 ± 19.14% for the BINAURAL BEATS whereas the ENVELOPE stimulation resulted in a mean increase of 70.09 ± 14.71% compared to the SHAM condition. After the linear mixed-effects model showed a significant effect of condition (see section above), post-hoc comparisons revealed a significant difference between the maximal Hilbert amplitude of the BINAURAL BEATS and ENVELOPE (p < 0.001). Furthermore, the increase in slow wave dynamics compared to SHAM was only significant within the first two seconds for the ENVELOPE stimulation and around 5 seconds for the BINAURAL BEATS condition (see Figure 2B). The topographical SWA effect of the auditory stimulation for every five seconds of the stimulation windows is illustrated in Figure 2C. We found no significant difference between ENVELOPE and SHAM condition for all 5 s windows after correction for multiple comparisons. On the other hand, BINAURAL BEATS stimulation significantly and globally enhanced SWA within the first five seconds. The relative change in SWA of both conditions compared to SHAM is shown in Figure A2A in the Appendix.

### Effects of auditory stimulation on sleep stage shifts and arousal

Considering the increase in power spectral density in almost the complete frequency spectrum, we wondered whether auditory stimulation increased the probability of wake epochs to occur and whether some conditions more likely woke up participants than others. We found a significant effect of condition (F_Cond_ (5,110) = 5.79, p_condition_ < 0.001) on the probability of wake epochs to occur. Post-hoc comparisons of the conditions revealed a significantly increased wake probability for BINAURAL BEATS condition tBinauralBeats(110) = 4.66, p < 0.001). All other conditions resulted in non-significant changes of the probability compared to the SHAM condition (all pCond > 0.1). In addition to sleep stage shifts towards wake epochs, we also compared the probability of automatically detected arousals occurring in a frontal (Fz), parietal (Pz), or occipital (Oz) electrode. We found no significant changes in either frontal, parietal or occipital detected arousals depending on condition (F_Cond_,Frontal(5,110) = 1.07, p = 0.38, F_Cond_,Parietal(5,110) = 1.10, p = 0.37, F_Cond_,Occipital(5,110) = 1.27, p = 0.28).

To further ensure that the observed brain oscillatory response is indeed not accompanied by any arousal, we analyzed the five seconds topographical response in the low alpha frequency band (8 - 10 Hz) which is shown in Figure A3 in the Appendix. For the DOWN and ISI1 stimulation, we observed significant increases in alpha during the first five seconds of stimulation, however, the effects were observed across all cortical regions and not only occipital which might point towards arousal (Pivik & Harman, 1995). Additionally, we found a significant increase in low alpha across all 5 s windows of the stimulation ON and stimulation OFF window for the BINAURAL BEATS, and this increase in low alpha became more prominent in occipital regions with time, indicating an arousal induced reaction. Because of a frequency overlap of alpha activity with low spindle activity, the increased alpha activity might potentially be caused by low spindle activity. Thus, we ran a spindle detection algorithm for all electrodes and calculated a spindle probability distribution our stimulation windows binned in two seconds. The probability distribution for slow (10 - 12 Hz) and fast (13 - 16 Hz) spindles for a frontal electrode (Fz), a centro-parietal electrode (CPz), and occipital electrode (Oz) are shown in Figure A4 in the Appendix. We found a significant increase in fast spindle probability especially in the frontal electrode for all conditions except for the ENVELOPE condition. Furthermore, we observed increased slow spindle probability in the frontal electrodes of all conditions. However, only for the BINAURAL BEATS, we observed significant increases in slow spindle activity also in the occipital electrode.

### Effects of sound volume on slow wave dynamics

Because we detected a large initial peak in the Hilbert slow wave amplitude in all stimulation conditions of Study1, we wondered whether we could reshape this initial peak by using sound volume modulation. Therefore, we employed again 1 Hz rhythmic stimulation at 45 dB (ISI1_High_) and a sound volume modulated condition where we increased the sound volume from 40 dB to 45 dB within five tones (ISI1_Mod_). As illustrated by the relative Hilbert amplitude shown in Figure 3A, the increasing sound volume of the ISI1_Mod_ significantly postponed the occurrence of the highest amplitude compared to the ISI1_High_ stimulation on an individual level by 4.39 s (t_Cond_(8) = 2.81, p = 0.02). Although the slow wave dynamics revealed significant differences between the two conditions in the beginning of the stimulation (Figure 3B), comparing the highest amplitude during the complete stimulation window of 39.37 ± 13.29% for the ISI1_High_ and 33.42 ± 7.42% for the ISI1_Mod_, showed no more significant difference between the two conditions (t_Cond_(8) = 0.64, p = 0.54).

**Figure 3:**
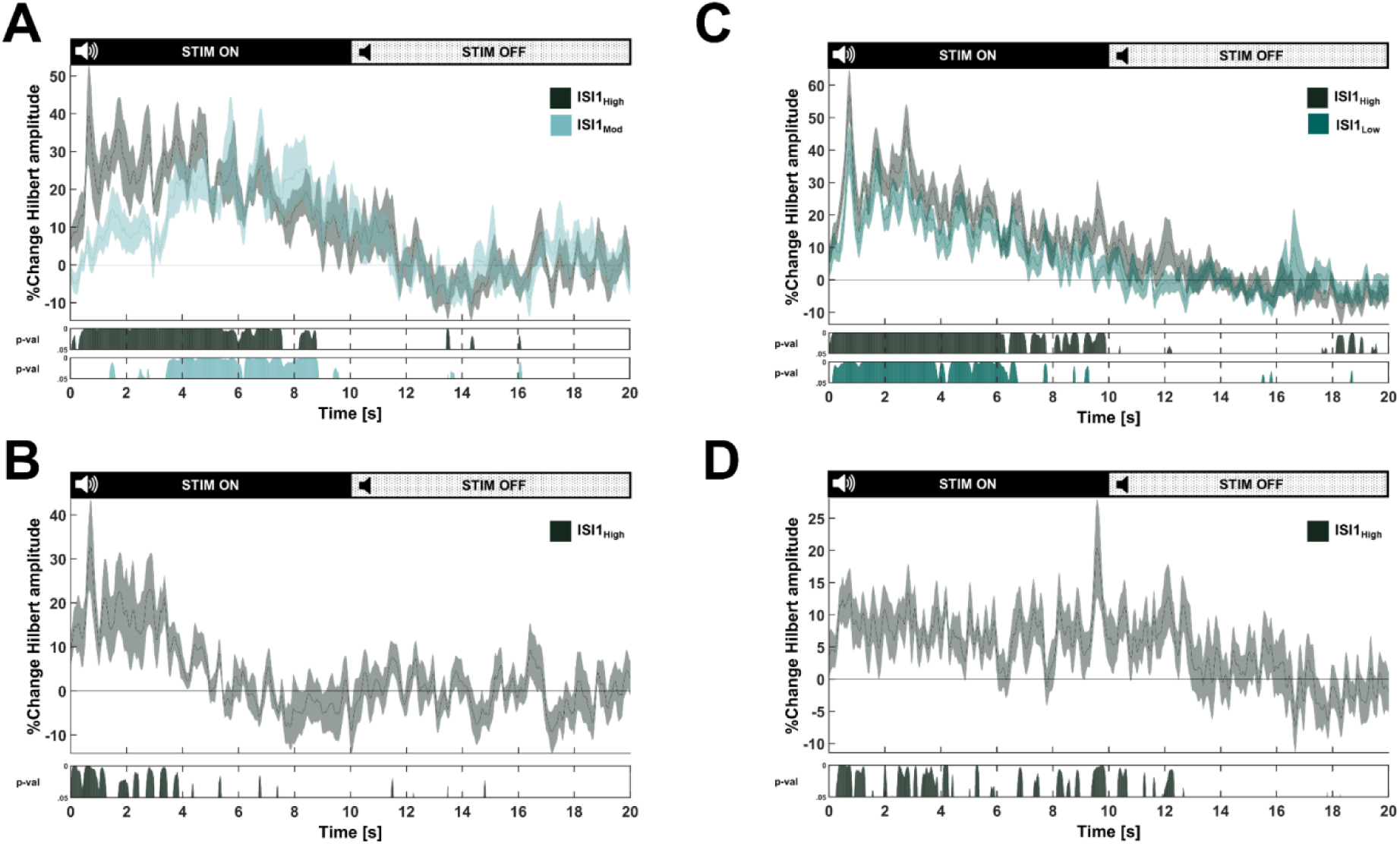
Results of 1 Hz rhythmic stimulation (ISI1) with different sound volumes on slow wave dynamics. A+C: Percentage change of Hilbert amplitude in the slow wave frequency band (0.5 – 2 Hz) for the conditions ISI1_High_ (45 dB), ISI1_Mod_ (40 - 45 dB) compared to SHAM (A) and ISI1_High_ (45 dB), ISI1_Low_ (42.5 dB) compared to SHAM (B), respectively. The horizontal line at zero represents no change compared to the SHAM condition. Below the plot the resulting p-values of the post-hoc comparison of linear mixed effects-models with condition as fixed factor and subject as random factor for each time point are shown. Dark green: ISI1_High_ to SHAM, bright blue: ISI1_Mod_ to SHAM (A), and dark green: ISI1_High_ to SHAM and turquoise: ISI1_Low_ to SHAM. B+D: Percentage change of Hilbert amplitude in the slow wave frequency band (0.5 – 2 Hz) for the condition ISI1_High_ compared to ISI1_Mod_ (C), and ISI1_High_ compared to ISI1_Low_ (B), respectively. The horizontal line at zero represents no change. Below the plot the resulting p-values of the post-hoc comparison of linear mixed effects-models with condition as fixed factor and subject as random factor for each time point are shown. Dark green: ISI1_High_ to ISI1_Mod_ (A), and dark green: ISI1_High_ to ISI1_Low_. Plots A + B: show n = 9 participants, and plot C + D: n = 19 participants.

Within Study3, we investigated whether 45 dB ISI1 (ISI1_High_) stimulation differed from 42.5 dB ISI1 stimulation (ISI1_Low_) to further elaborate on the effects of sound volume on auditory slow wave enhancement and whether there might be an accumulated dose-dependency effect. Figure 3C illustrates the relative Hilbert response of the ISI1_High_, the ISI1_Low_, compared to the SHAM showing significant differences in the Hilbert amplitude across the stimulation ON window between the conditions and SHAM. The highest amplitude at the location of the maximum of the Hilbert amplitude of the ISI1_High_ was 56.72 ± 8.05% and 39.75% ± 7.45% for the ISI1_Low_. A paired t-test revealed a significant difference (t_Cond_(18) = 3.42, p = 0.003) between the maximal amplitude of the ISI1_High_ and ISI1_Low_. As illustrated in Figure 3D which shows the relative difference in the Hilbert amplitude between the ISI1_High_ and ISI1_Low_, there was an increase for the ISI1_High_ across the complete stimulation ON window.

### Effects of time of the night on the efficacy of auditory stimulations

Because previous research using auditory stimulation to modulate sleep slow waves often only stimulated for a certain time (Besedovsky et al., 2017; Hong-Viet V. Ngo, Martinetz, et al., 2013) we wanted to investigate whether we observe a different effect on the brain depending on the time of stimulation. Thus, we divided the night into first four hours of stimulation, starting from the first stimulation ON window, and remaining hours of stimulation. The number of stimulation windows during early and late nights are shown in the Appendix in Table A2. Paired two-sided t-tests between the number of stimulation windows revealed a significant larger number of stimulation windows during early nights for all studies separately (Study1: t_Time(22)_ = 4.13, p < 0.001; Study2: t_Time(8)_ = 7.30, p < 0.001; Study3: t_Time(18)_ = 6.76, p < 0.001). Next, we wanted to check whether the number of stimulations within a stimulation ON window differed for the early or late night. A paired two-sided t-test revealed a significantly higher number of stimulations occurring during the stimulation windows in the early nights of the UP condition (t_Time(20)_ = 2.93, p = 0.008). On the other hand, we did not detected a significant difference in number of stimulation for the DOWN condition (t_Time(20)_ = 1.58, p = 0.13).

As a next step, we also calculated the SWA for 5 s windows of the stimulation window. As illustrated in Figure 4, the SWA enhancement during the first five seconds is present during early and late nights. However, we detected only a significant SWA enhancement for the second five seconds during early nights. Furthermore, as shown in Figure A5 in the Appendix, if calculating the SWA across the complete stimulation window, we observed a significant effect of condition only for the early nights (F_Cond_(5,110) = 9.51, p < 0.001), and post-hoc comparisons showed a significant SWA enhancement for all conditions except the ENVELOPE condition (p < 0.05, except p_ENVELOPE_ = 0.89). However, comparing the SWA of the late night, the linear mixed-effects model was not significant for condition (F_Cond_(5,101.12) = 1.78, p = 0.12). Furthermore, we also calculated the Hilbert amplitude response in the low slow wave frequency bands (0.5 - 2 Hz) for early and late nights for the ISI1_High_ condition and SHAM condition only. These results are illustrated in Figure A5B in the Appendix. The plot matches with our SWA enhancement by that the Hilbert amplitude of the ISI1_High_ seems to be enhanced for a longer time compared to the SHAM for the early night, and this enhancement is less pronounced during late nights.

**Figure 4:**
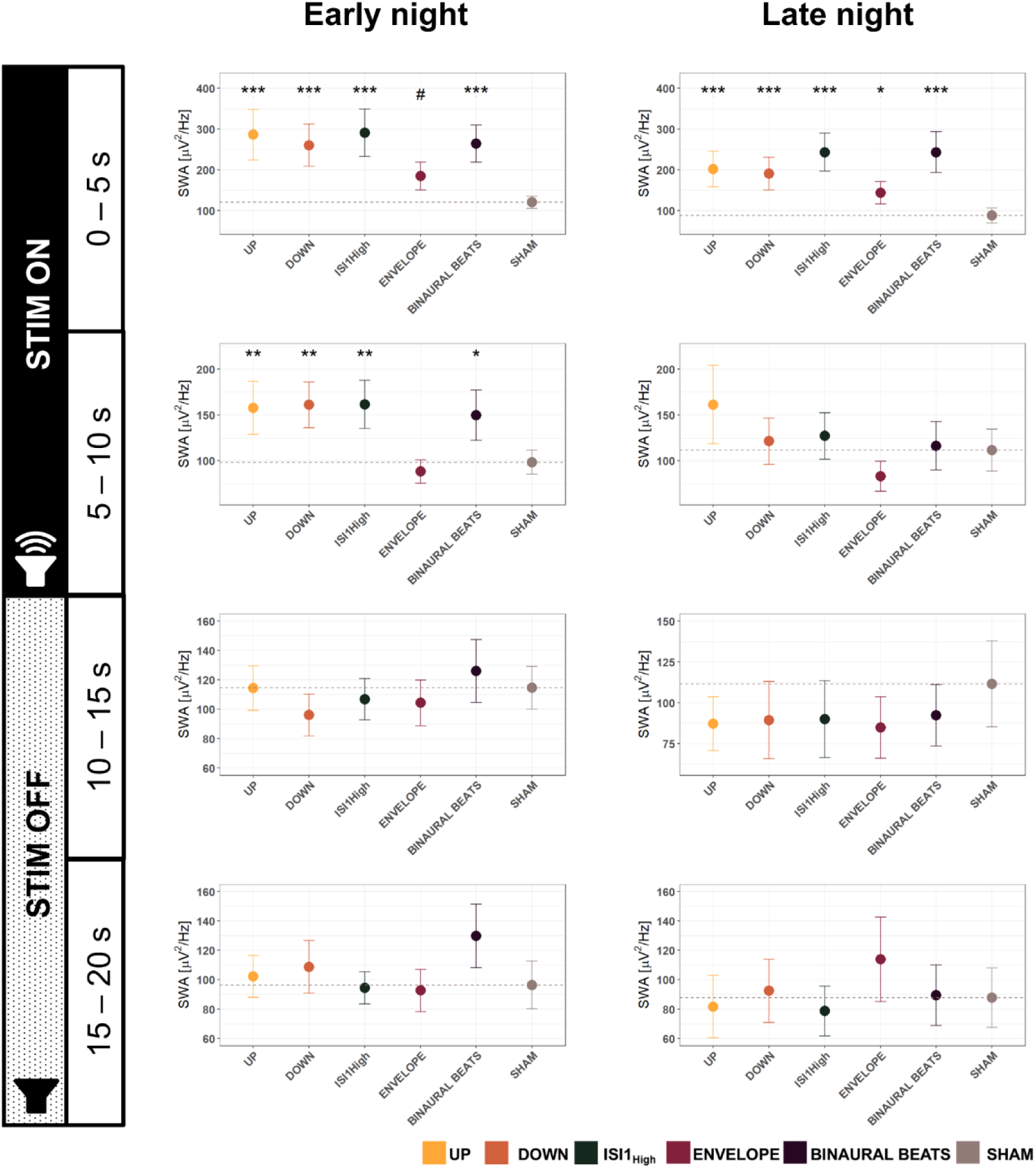
Effects of time of the night on slow wave activity (SWA; 0.5 – 2 Hz). Early night shows SWA data of the Fz electrode for all stimulation conditions (UP, DOWN, ISI1_High_, ENVELOPE, BINAURAL BEATS, SHAM) for the first four hours after the first stimulation window started. Late night represents the remaining hours of stimulations. We employed linear mixed effect models for all five seconds of the stimulation windows entering the stimulation condition as fixed effect and subject as random factor. All post-hoc p-values represent the significance compared to the SHAM condition and have been adjusted for multiple comparisons using the Hochberg correction. ***: p < 0.001, **: p < 0.01, *: p < 0.05, #: 0.05 < p < 0.1. All data is shown for n = 23 participants.

As a last step we investigated whether time of the night influences the probability of wake epochs to occur upon stimulation. Because we only observed significant effects of the BINAURAL BEATS condition on wake probability, we employed a linear mixed-effects model with the interaction of time of the night and condition as fixed factors with only the SHAM and BINAURAL BEATS conditions, which was not significant (F_Cond x Time_(1,64.061) = 0.028, p = 0.87). Running a linear mixed-effects model for the early and late night separately showed a significant effect of condition (F_Cond,Early_(1,22) = 10.902, p = 0.003) and an increase in the probability by 8.96% for the BINAURAL BEATS. The same model showed trend level (F_Cond,Late_(1,20.942) = 3.57, p = 0.07) for the late night, with an increase of the wake probability by 8.95%.

### Effects of auditory stimulation on cardiovascular dynamics

Because the brain and the body are not uncoupled during sleep and the ANS represents a key pathway between the brain and body, we first wanted to investigate whether there are immediate, dynamical changes in IHR during the stimulation windows. To overcome the general decline in heart rate occurring with the course of the night (Cajochen et al., 1994), we calculated a relative IHR compared to the mean heart rate of the closest SHAM stimulation window if this SHAM window occurred within five minutes. As illustrated in Figures 5A and B, we observe a slight acceleration in the IHR during the first four seconds of all auditory stimulation conditions followed by a longer deceleration of IHR compared to the SHAM condition. To quantify these dynamic effects, we calculated the mean RR interval (Note: inverse of heart rate, the longer the RR interval, the slower the heart rate), the longest and shortest RR interval of each stimulation window, the difference between the shortest and longest RR interval of each stimulation ONOFF together window, and the short time HRV indices RMSSD and SDNN. As shown in Figure 3C, we found a trend-level effect of the stimulation conditions on the prolongation of the mean RR interval (F_Cond_ (5,110) = 2.05, p = 0.078), with post-hoc comparisons indicating significance (p < 0.05) only for the DOWN and ISI1_High_ condition. Furthermore, we detected a significant influence of conditions for the longest RR interval (F_Cond_ (5,110) = 6.10, p < 0.001), and post-hoc comparisons showed a significant prolongation for all conditions compared to SHAM (all p < 0.01). Analyzing the shortest RR interval in the stimulation window, we detected a significant influence of condition (F_Cond_ (5,110) = 5.69, p < 0.001), and post-hoc comparisons between conditions revealed that, only BINAURAL BEATS significantly shortened the shortest RR interval in the stimulation window compared to SHAM (p < 0.001). To quantify the ANS influence on the heart, we compared two HRV indices: First, we found a significant effect of condition on RMSSD (F_Cond_ (5,110) = 5.63, p < 0.001), and post-hoc comparisons between the conditions showed a significant increase in RMSSD compared to SHAM for the UP, DOWN, and BINAURAL BEATS (all p < 0.05), and a trend for ISI1_High_ (p = 0.0504). Moreover, we detected a significant influence of condition on SDNN (F_Cond_ (5,110) = 7.72, p < 0.001). Post-hoc comparisons revealed a significant increase for UP, DOWN, ISI1_High_, and BINAURAL BEATS (p < 0.05), whereas the effect of ENVELOPE remained on trend-level (p = 0.0521).

**Figure 5:**
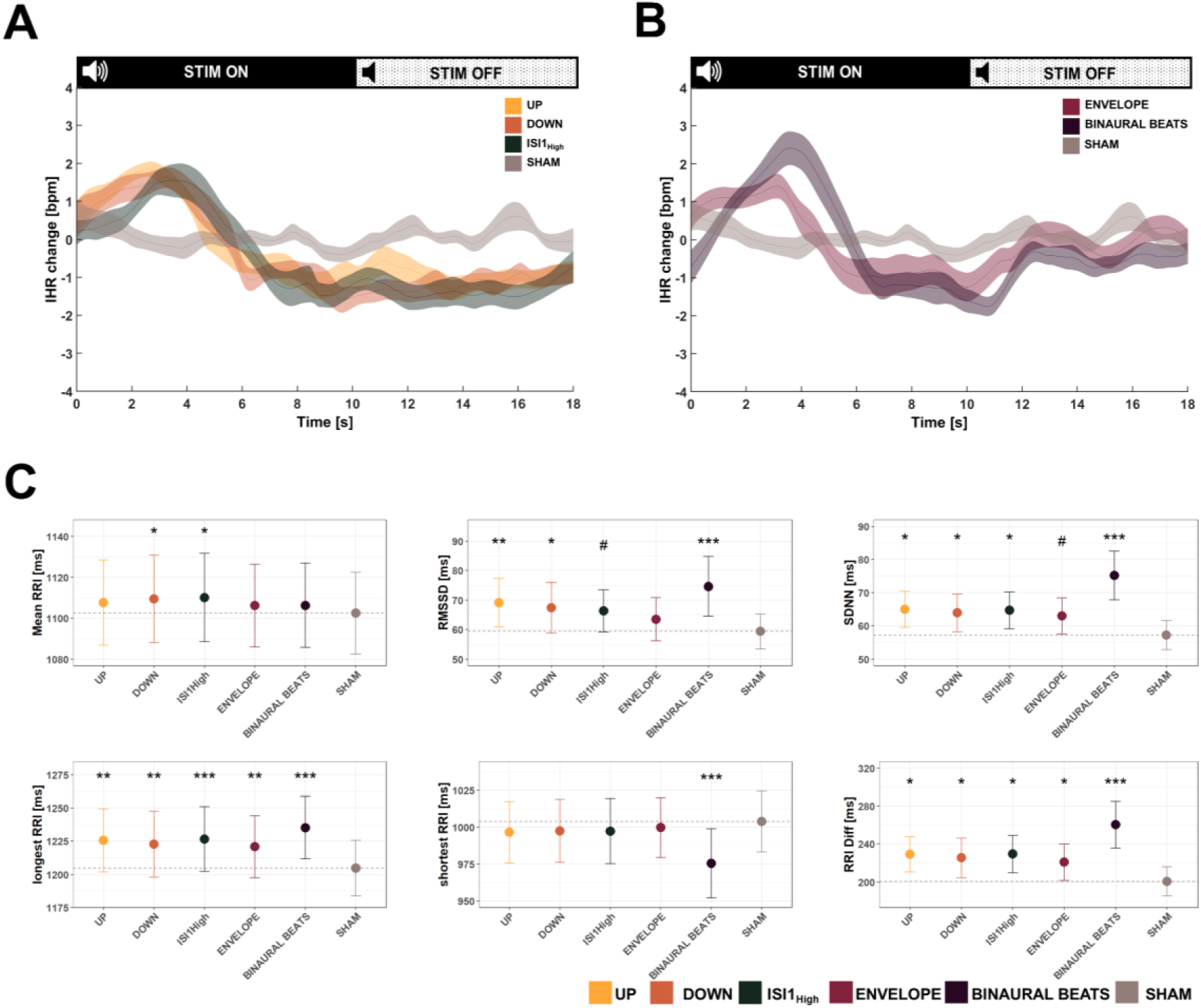
Cardiovascular response to auditory stimulation. A + B: Change in instantaneous heart rate (IHR) during stimulation window for UP, DOWN, IS1High and SHAM stimulation (A), or ENVELOPE, BINAURAL BEATS and SHAM (B) compared to the mean of the closest SHAM window, if the closest window was within five minutes. IHR is presented as mean ± standard error of the mean. C: Heart rate (variability) features for stimulation windows for the conditions UP, DOWN, ISI1High, ENVELOPE, BINAURAL BEATS, and SHAM stimulation. All data is presented as mean ± standard error of the mean. RRI: Interval between two normal heart beats, RMSSD: Root mean square of successive differences between normal heart beats. SDNN: Standard deviation of normal heart beats. Longest RRI: Duration of the longest interval between two consecutive heart beats in a stimulation window. Shortest RRI: Duration of the shortest interval between two consecutive heart beats in a stimulation window: RRI Diff: Difference between the longest and shortest interval within a single stimulation window. Note that the RRI are inversely correlated with heart rate, thus, the longer the RRI, the lower the heart rate. Significance levels of the linear mixed-effects model with condition as independent factor and subject as random factor are shown for each cardiovascular feature in the plot and corrected for multiple comparisons using the Hochberg method. ***: p < 0.001, **: p < 0.01, *: p < 0.05, #: 0.05 < p < 0.1. Data is shown for n = 23 participants.

**Figure 6:**
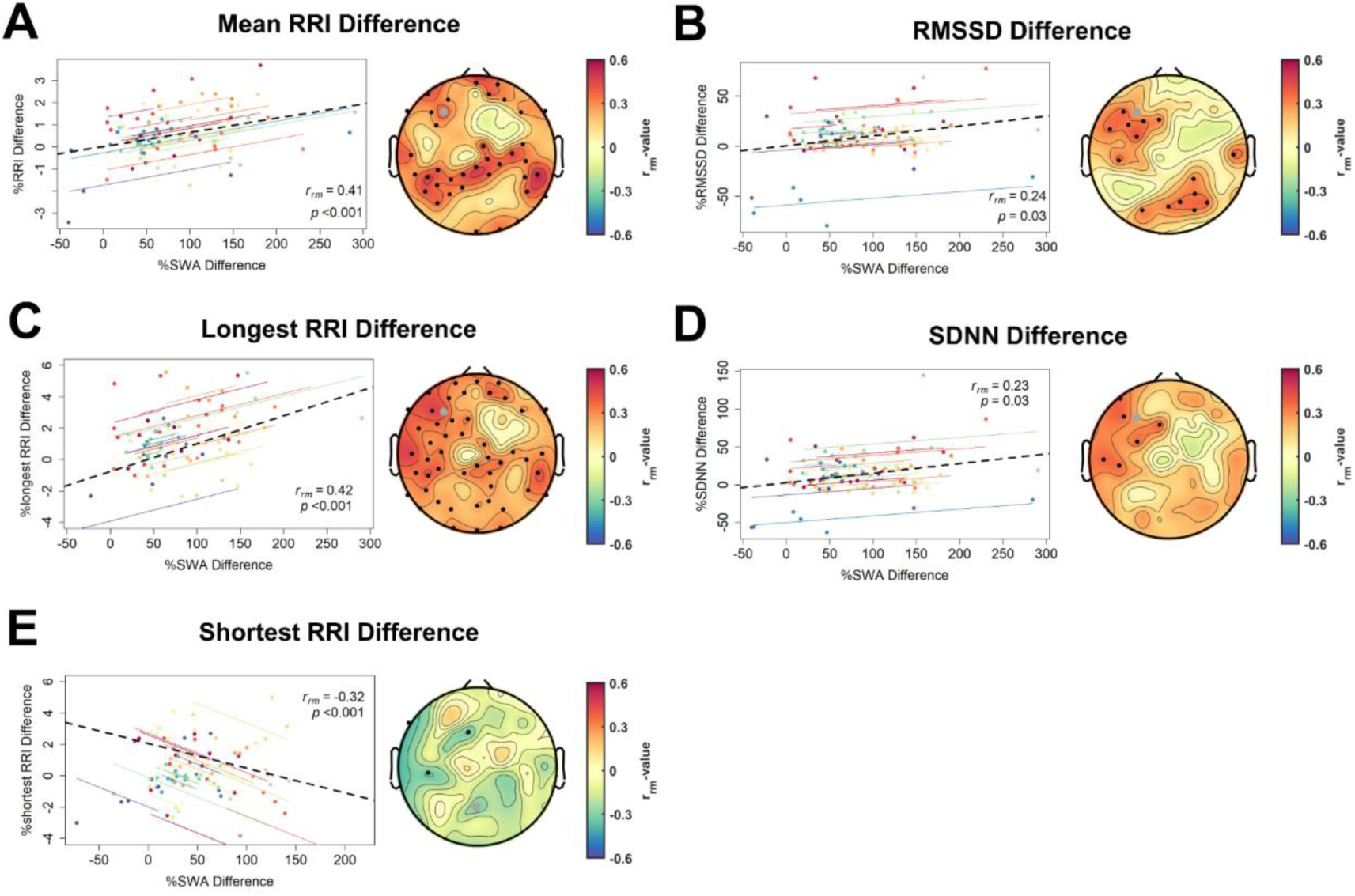
Repeated measures correlations between percentage change for heart rate (variability; HR(V)) features and percentage change of slow wave activity (0.5 – 2 Hz) in the first five seconds of the stimulation ON window. For every participant (n = 22) and every condition a mean value for each variable was calculated. We excluded one participant because not all conditions had ECG measurements of high enough qualities. Topoplots show r values and significant electrodes (p < 0.05) are marked with black dots. We corrected the p-values for multiple comparisons using false discovery rate. The repeated measures correlation is plotted for the example electrode F3 (marked with a grey point in the topoplots) if not otherwise mentioned, and the respective p-values and rrm values are shown in the plots. A: Mean difference of consecutive heart beats (RRI). B: Difference in the root mean square of successive difference of normal heart beats (RMSSD). C: Differences of the longest RRI, D: Difference in the standard deviation of differences between normal heartbeats (SDNN). E: Percentage of difference of the shortest RRI correlated with percentage change in SWA for electrode Pz.

As a next step, we investigated whether the auditory induced SWA enhancement correlated with the above-mentioned cardiovascular changes. Thus, we conducted a repeated measures correlation between percentage change of every stimulation condition compared to SHAM. For the cardiovascular changes, we used the complete stimulation window indices whereas we took the SWA of the first five seconds of the stimulation ON window as we suspected those effects to be the driver of the cardiovascular changes considering the time course of the IHR. As illustrated in Figure 4, we found a significant, strong, and positive correlation between the difference in SWA and the difference in mean RR interval, and the difference in the longest RR intervals, particularly for frontal and parietal electrode clusters. Furthermore, we found some frontal and occipital clusters showing a significant positive correlation of the change in SWA with RMSSD and SDNN. Only a few electrodes showed significant negative correlations with the shortest RR interval, indicating that the higher the change in SWA the shorter the shortest RR interval. Furthermore, we also correlated the cardiovascular parameters with changes in the low alpha (8 - 10 Hz) frequency band, which is illustrated in Figure A6 in the Appendix. We found SDNN to be positively correlated with an increase in low alpha across a cluster of right frontal, central, and parietal electrodes.

## Discussion

In this paper, we compared various widely used auditory stimulation conditions with a SHAM condition within a single sleep period and found strong and global enhancements of SWA of all stimulation conditions except the ENVELOPE condition. Furthermore, the increase in slow wave dynamics within stimulation windows was most pronounced during the beginning of stimulation and decreased with the duration of stimulation. Additionally, the time course of the increase in SWA can be altered by modulating the sound volume of the stimulation. We also showed that auditory stimulation applied during the second half of the night had a decreased efficiency of enhancing SWA. Moreover, we provide first evidence on the effects of auditory slow wave modulation on cardiovascular dynamics. We showed that auditory stimulation modulates the course of the IHR during the stimulation windows and changed cardiovascular parameters towards more resting cardiovascular conditions. Importantly, the extent of the cardiovascular change was correlated with the change of SWA, presumably indicating slow waves to be the driver of these changes. Altogether, we presented a novel stimulation protocol allowing resource-efficient comparisons between different auditory stimulation conditions on dynamic changes on slow wave and cardiovascular levels.

### Auditory stimulation globally enhances SWA

We found a strong and global enhancement of SWA for all tested stimulation conditions except for the ENVELOPE condition. This enhancement was independent of target phase of the auditory stimulation. Contrary to previous studies (Fattinger et al., 2017; Hong-Viet V. Ngo, Martinetz, et al., 2013) we did not observe a decrease in SWA and slow wave dynamics of the down-phase stimulation compared to the SHAM condition, however, we even observed significant increases of SWA on a global cortical level. Comparing our results to the stimulation design of Fattinger and colleagues, we did not stimulate every down-phase of a slow wave, but only slow waves in the 10 s stimulation ON window of the DOWN condition. The slow wave decreasing effects may accumulate over time of stimulation and our usage of various auditory stimulation conditions might have interfered with the slow wave diminishing effects. Interestingly, we observed significant decreases in high SWA (2.25 - 4 Hz) over central areas during the stimulation OFF window, overlapping the targeted premotor cortex of Fattinger’s study (Fattinger et al., 2017). However, these decreases have been observed for all reported stimulation conditions of Study1 and are thus not dependent on the target phase of the stimulation. We, therefore, conclude that global SWA decreases are not possible using auditory slow wave modulation. Another possible explanation why we found contrary results to Fattinger and colleagues could be that they used a stimulation electrode located above the sensory-motor cortex and the stimulation only served to perturb slow waves on a local cortical level. We on the other hand targeted slow waves at the frequent prefrontal origin of slow waves from where the slow waves can propagate across the complete cortex (Massimini et al., 2004). Thus, we cannot exclude that the mechanism behind local and global slow wave down modulation might be different. Lastly, high-volume sounds have been previously applied to strongly perturb slow waves, but this stimulation also elicited arousals (Landsness et al., 2009; Tasali et al., 2008), and increased wake (Aeschbach et al., 2008). Thus, sound volume may be an important factor to consider for perturbing slow waves and further studies are needed to investigate the underlying mechanisms on global (e.g., frontal) and local slow wave modulation.

### Effect on SWA decreases with duration of auditory stimulation

We observed the strongest increases in SWA and slow wave dynamics during the beginning of stimulation and this enhancement decreased with time of stimulation, although for all conditions except the UP and DOWN phase targeting condition, auditory stimulation was present during the complete stimulation ON window. In line with this finding is another study of Ngo and colleagues (H.-V. V Ngo et al., 2015) where they investigated the efficiency of a two stimulation protocol to an ongoing stimulation protocol in enhancing slow waves and subsequent memory consolidation. They found no difference in the enhancement of slow waves or induction of trains of slow waves between the two conditions, concluding that there is a mechanism at play preventing the brain from hypersynchrony caused by slow wave enhancement. Furthermore, Lustenberger and colleagues (under review) showed that the overall effect of continuous auditory stimulation on SWA compared to windowed ONOFF stimulation did not differ between the conditions, although around double of the stimulations were applied in the continuous approach. Both studies indicate that breaks between stimulations favor subsequent auditory stimulation induced slow wave enhancements.

As already proposed by Bellesi et al. (Bellesi et al., 2014), auditory stimulation during sleep may activate the non-lemniscal auditory processing pathway, and the target dorsal and caudo-medial medial geniculate body of the thalamus (part of the auditory thalamus) that activates the cortex to changing acoustic environment (Kraus et al., 1994). This global activation of the cortex is reflected by large amplitude and steep slow waves reflecting an efficient neuronal synchronization process that could be termed as K-complex or Type 1 slow wave (Bernardi et al., 2018). K-complexes commonly arise as a reaction to sensory stimulation and serve to protect sleep continuity (e.g., reviewed in (Parrino & Vaudano, 2018), (Bernardi et al., 2018)). However, K-complexes can be associated with arousal reactions, indicating activation of the activating reticular ascending system (ARAS). The locus coeruleus (LC) as the main source of norepinephrine in the brain is a key player to modulate arousal levels (Carter et al., 2010). Beyond modulating arousal levels, the LC may also have inhibitory effects on a sensory level (e.g., reviewed in (McBurney-Lin et al., 2019)), and this inhibitory effect caused by the LC has also been reported in the auditory system (Foote & Morrison, 1987). Therefore, the decrease of the response to auditory stimulation might not only be caused by cortical or cortico-thalamic processes but already start with an adaptation of the response on the auditory sensor level or at the level of the LC, which may provide further evidence of the importance of breaks being important for efficient auditory stimulation. Therefore, we strongly propose to include breaks (e.g., using a windowed stimulation approach) for efficient auditory slow wave enhancement. Nevertheless, future research is needed to further elaborate on the reasons for the decline in efficacy of auditory stimulation with the duration of stimulation.

### Auditory slow wave enhancement is not accompanied by arousals

Given the previous indications that the effect of the auditory slow wave modulation might be arousal driven as the shape of the first increase in the Hilbert amplitude strongly resembles a K-complex, which was proposed to evoke sleep spindles (Steriade, 2006), we wanted to investigate whether the observed SWA increase might be accompanied by arousals and therefore, a non-specific increase across the complete EEG power spectrum. Particularly an increase in occipital low alpha activity could indicate an auditory induced arousal. Interestingly, we only found binaural beats to increase the alpha activity of the stimulation ON and OFF window, and this increase became more occipitally focused with duration of the stimulation OFF window. However, because spindles and alpha activity show overlapping EEG frequency bands, it is difficult to distinguish those just based on EEG power. Analyzing a spindle probability distribution for slow and fast spindles to account for the fact that there is a spindle gradient showing slower frontal and faster centro-parietal spindles (Andrillon et al., 2011), we found increases in slow and fast frontal spindle probability compared to SHAM. Furthermore, there were only slight differences in fast occipital spindles for some of the conditions. However, only binaural beats show significant increases in slow occipital spindles. Taken together with the significant increase of the probability of a wake epoch to co-occur with BINAURAL BEATS stimulation, we assume that only BINAURAL BEATS stimulation induced arousals. A possible explanation why BINAURAL BEATS more likely woke or aroused participants could be that although we used 45 dB for all conditions of Study1, BINAURAL BEATS was the only one condition playing a continuous, pure-frequency sound for 10 s, most other conditions only used short, 50 ms bursts of 45 dB pink noise.

### Stimulations at the end of the night are less efficient in enhancing SWA

Many of the previously published studies only applied auditory stimulations for a certain amount of time during the beginning of the night (Besedovsky et al., 2017; Hong-Viet V. Ngo, Martinetz, et al., 2013; H.-V. V Ngo et al., 2015; Weigenand et al., 2016) whereas others stimulated during the complete night (Grimaldi et al., 2019; Leminen et al., 2017). Thus, we wanted to elucidate whether there is a different SWA response to auditory stimulation depending on the time of the night. Our results revealed that stimulation is more efficient during the first four hours of stimulation compared to the remaining hours of stimulation. Interestingly, there was no significant slow wave enhancement during the late night when considering the ON and OFF window together. Additionally, there was a shorter increase in slow wave dynamics during late nights. This less efficient stimulation towards the end of the night might be caused by generally decreased SWA (D.-J. J. Dijk, 2009) and less occurring high amplitude slow waves (Riedner et al., 2007) leading to less slow waves that could be stimulated. However, only the UP and DOWN condition depended on the presence of slow waves within the stimulation ON window, whereas for the ISI1_High_ condition, the same number of stimulations within the stimulation ON was applied in early and late nights. Therefore, the presence of slow waves per se cannot be the cause for the decrease in stimulation efficacy. Nevertheless, the decrease in sleep pressure reflected by decreased SWA (Achermann et al., 1993) accompanied by a reduction in synaptic strength (Esser et al., 2007), might also indicate slow wave saturation effects, meaning that with duration of the night, the ability of the system to enhance slow waves might be limited. However, the generation of large subcortico-cortical neuronal synchronization reflected by K-complexes is not perturbed with duration of the night and K-complexes even tend to become larger during later parts of the night (Bernardi et al., 2018). In contrast, slow wave amplitude and slow wave slopes are generally decreased in later parts of the night (Riedner et al., 2007) and thus, only the cortico-cortical slow waves might be influenced by those possible saturation effects. Altogether, we provide further evidence to restrict slow wave sleep modulation to NREM sleep during the early times of the night, where stable NREM sleep is dominating, and therefore provide efficient slow wave modulation. Nevertheless, establishing the mechanisms that explain the change of response to auditory stimulation across the sleep period need further investigation.

### Auditory slow wave enhancement alters cardiovascular dynamics, and those changes are correlated to changes in SWA

We wanted to elucidate whether we detect changes in cardiovascular dynamics time-locked to the auditory stimulation windows. Considering the course of the IHR, we observe clear differences between the auditory stimulation conditions and SHAM. The time course might indicate at first sight that there was a first acceleration of the heart rate that would indicate an arousal on a cardiovascular level. This initial visually observed increase in IHR could indicate an increase in sympathetic activity at first sight. However, sympathetic modulation needs more than five seconds until it changes cardiovascular functions (Nunan et al., 2010). On the other hand, parasympathetic activity rapidly changes the heart rate, therefore, being the main contributor to beat-to-beat variations of the heart rate (Shaffer & Ginsberg, 2017). Thus, the initial IHR course may rather be the outcome of decreased parasympathetic activation with the beginning of the stimulation. This might be an undesired cardiovascular response, however, we found no statistical evidence of a shortening of the shortest RR interval for all conditions, except the BINAURAL BEATS. Altogether, future studies should not only consider that the effects of auditory stimulation are not limited to the brain only, but that they might even entail unfavorable changes on the cardiovascular system as seen by the BINAURAL BEATS stimulation.

The initial acceleration of IHR was followed by a longer-lasting period of deceleration as reflected by a prolongation of the longest RR interval for all conditions, and even a significant prolongation of the mean RR interval for the DOWN and ISI1_High_ stimulation, indicates a significant slowing of the heart rate. This change in cardiovascular dynamics was also reflected by the underlying change in cardiac autonomic regulation as reflected by the HRV changes. The observed increased SDNN for all conditions (except only trend level for ENVELOPE) points towards generally increased autonomic activation (Shaffer & Ginsberg, 2017) and the significantly increased RMSSD for the UP, DOWN, BINAURAL BEATS, and a trend for ISI1_High_ indicates increased parasympathetic activation. Our finding is in line with correlational evidence showing increased HRV parameters with increased SWA (Brandenberger et al., 2001; Jurysta et al., 2003; Otzenberger et al., 1998), and a previous study of Grimaldi demonstrating effects of auditory stimulation on five-minutes HRV measurements (Grimaldi et al., 2019). However, the HRV results should always be interpreted with caution, as those measurements only indirectly reflect underlying ANS activity. But by taking into account the IHR or heart rate over the complete stimulation window, we can verify whether the observed HRV changes also correspond with the (I)HR as outcome of the cardiac autonomic regulation reflected by HRV. As for instance seen with BINAURAL BEATS, we observed a significant increase in RMSSD, alongside with a significant shortening of the shortest RR interval. Together with the higher probability of inducing arousal-alike brain traits, we suggest interpreting the increased RMSSD with caution. Nevertheless, for all other conditions, the detected changes in HRV match with the IHR and RR interval findings.

As a next step, we showed that the auditory evoked SWA enhancements directly translates to temporally coupled changes in the cardiac autonomic regulation, pointing towards more restful cardiovascular conditions when slow waves are prevailing. Although the induced changes on the heart are rather small (e.g., prolongation of the longest RRI by around 20 ms), we also need to consider the short time scale of only 10 seconds stimulation for this cardiovascular change to occur. Because we included all conditions in this correlation and not all conditions caused a similar effect on SWA, our results may indicate the more SWA we induce, the stronger the slowing of the heart. The effect of increased SWA over a certain time may also accumulate to stronger slowing of the heart, as it can be observed in normal sleep as well. Therefore, effects of the auditory slow wave enhancement may accumulate over time to increase parasympathetic predominance and lower the heart rate more substantially. Autonomic imbalance caused by sympathetic overactivity or parasympathetic underactivity has been related to many unfavorable effects on the brain and body (e.g., it is a common mechanism of a majority of risks factors to develop cardiovascular diseases (Thayer et al., 2010), the number one cause of death in the world (WHO, 2016)). Additionally, as aging, which is the number one risk factor for developing cardiovascular diseases (Landolt et al., 1996), is accompanied with a significant decrease in SWA (Lakatta, 2003), auditory slow wave enhancement with optimal settings may be a mean to counteract those unfavorable changes. However, applying one optimal auditory stimulation condition is needed to elucidate possible effects accumulated over stimulation time on heart rate and whether there are additional cardiovascular benefits for instance lowered blood pressure, that could potentially be related to slow waves.

### Limitations

Because we wanted to maximize the number of stimulation windows but still reliably compare HRV features, we calculated HRV for ∼20 s windows. This is a very short time based on official guidelines provided by the Task Force of The European Society of Cardiology and The North American Society of Pacing and Electrophysiology (Schlosser, 1996). Thus, we cannot generalize our HRV findings to clinical HRV analysis as e.g., 24h HRV recordings because the much longer time duration does not allow for this comparison. However, to overcome this issue, we analyzed the time-domain HRV measurements RMSSD and SDNN and compare the stimulation effects between the conditions only. Additionally, RMSSD was already shown to highly correlate in 10 s recordings compared to standard five minutes measurements (Thong et al., 2003). Furthermore, the here presented results are intended to be used as guidelines for stimulation effects during the stimulation window itself. Therefore, we cannot draw any conclusion whether the same effects will be obtained if any of the here reported stimulation condition is applied as single condition across a complete night. Such a study would also allow to test behavioral and functional effects of the stimulations, as for instance overnight memory consolidation or ANS testing.

## Conclusion

Here, we aimed to provide a framework for comparing auditory stimulation conditions in a novel within-night protocol on brain and cardiovascular responses. First, we found no evidence that phase of stimulation influences SWA enhancement and were thus not able to replicate previous research showing decreases in SWA upon down-phase stimulation. Additionally, we showed that stimulation induced SWA enhancement is most pronounced at the beginning of a stimulation window and decreases with the time of stimulation. Moreover, we also demonstrated that by changing the volume of stimulation the SWA enhancement can be modulated and that stimulation towards the end of the night gets less successful in enhancing SWA compared to earlier times of the night. However, the underlying causes for the decline in the SWA enhancement with the duration of stimulation and with time of the night remain to be elucidated. Lastly, we provide first evidence that auditory stimulation directly affects the heart on a stimulation window level pointing towards more restful cardiovascular conditions as indicated by a prolongation of the longest RR interval, reflecting a slowing of the heart rate. Most importantly, these cardiovascular changes are correlated with auditory stimulation induced SWA enhancement indicating increased restorative functions when slow waves are prevailing.

## Data availability

Source data required to reproduce the main figures and main conclusions of the manuscript will be made available in the future.

## Code availability

Customized codes used to create the main figures and main conclusions in the manuscript will be made available in the future.

## Acknowledgements

All authors thank their trainees, collaborators, and mentors for the expiring exchanges and dialogues on the thematic of this article. The corresponding author Caroline Lustenberger is supported by the Swiss National Science Foundation (PZ00P3_179795) and this work was conducted as part of the SleepLoop Flagship of Hochschulmedizin Zürich. We thank Dr. M. Laura Ferster for sharing her expertise for the implementation of the automatic stimulation protocol. SleepLoop consortium members provided uncountable discussions and feedback. We are grateful to the Alumni Association of ETH Zurich for their support in participant recruitment for this study. We also thank Prof. Dr. Rafael Polania for his continuous support and feedback during the study.

## Author Contributions

St.H.: Conceptualization, methodology, software, formal analysis, investigation, data curation, writing - original draft, writing - review & editing, visualisation, project administration. M.C.D: Investigation, validation, writing - review & editing. Si.H.: formal analysis, investigation. L.K: investigation. F.S: Investigation. R.S: Formal analysis, investigation. F.A: Investigation. A.T: Formal analysis, investigation. C.S: Conceptualisation, writing - review & editing. R.H: Conceptualisation, methodology, writing - review & editing. N.W: Conceptualisation, supervision, resources, writing - review & editing. C.L: Conceptualisation, methodology, validation, resources, writing - original draft. Writing - review & editing, supervision, project administration, funding acquisition. All co-authors reviewed and edited the manuscript, before approving its submission.

## Conflict of Interest Statement

C.L. is a member of the Scientific Advisory Board of Emma Sleep GmbH, which is not related to this work. R.H. is a founder and shareholder of tosoo AG.

## Notes

### Summary of Updates

We updated the acknowledgement.

## Bibliography

Achermann, P., Dijk, D. J., Brunner, D. P., & Borbély, A. A. (1993). A model of human sleep homeostasis based on EEG slow-wave activity: Quantitative comparison of data and simulations. Brain Research Bulletin, 31(1–2), 97–113. https://doi.org/10.1016/0361-9230(93)90016-5

Aeschbach, D., Cutler, A. J., & Ronda, J. M. (2008). A role for non-rapid-eye-movement sleep homeostasis in perceptual learning. Journal of Neuroscience, 28(11), 2766–2772. https://doi.org/10.1523/JNEUROSCI.5548-07.2008

Alvarez-Estevez, D., & Fernández-Varela, I. (2019). Large-scale validation of an automatic EEG arousal detection algorithm using different heterogeneous databases. Sleep Medicine, 57, 6–14.

Andrillon, T., Nir, Y., Staba, R. J., Ferrarelli, F., Cirelli, C., Tononi, G., & Fried, I. (2011). Sleep Spindles in Humans: Insights from Intracranial EEG and Unit Recordings. Journal of Neuroscience, 31(49), 17821–17834. https://doi.org/10.1523/JNEUROSCI.2604-11.2011

Bakdash, J. Z., Marusich, L. R., & Marusich, M. L. R. (2016). Package ‘rmcorr.’

Bates, D., Mächler, M., Bolker, B. M., & Walker, S. C. (2015). Fitting linear mixed-effects models using lme4. Journal of Statistical Software. https://doi.org/10.18637/jss.v067.i01

Behar, J. A., Rosenberg, A. A., Weiser-Bitoun, I., Shemla, O., Alexandrovich, A., Konyukhov, E., & Yaniv, Y. (2018). PhysioZoo: A novel open access platform for heart rate variability analysis of mammalian electrocardiographic data. Frontiers in Physiology, 9(OCT), 1–14. https://doi.org/10.3389/fphys.2018.01390

Bellesi, M., Riedner, B. A., Garcia-Molina, G. N., Cirelli, C., & Tononi, G. (2014). Enhancement of sleep slow waves: Underlying mechanisms and practical consequences. Frontiers in Systems Neuroscience, 8(October), 1–17. https://doi.org/10.3389/fnsys.2014.00208

Benjamini, Y., & Hochberg, Y. (1995). Controlling the False Discovery Rate: A Practical and Powerful Approach to Multiple Testing. Journal of the Royal Statistical Society: Series B (Methodological), 57(1), 289–300. https://doi.org/10.1111/J.2517-6161.1995.TB02031.X

Bernardi, G., Siclari, F., Handjaras, G., Riedner, B. A., & Tononi, G. (2018). Local and widespread slow waves in stable NREM sleep: Evidence for distinct regulation mechanisms. Frontiers in Human Neuroscience, 12(June), 1–13. https://doi.org/10.3389/fnhum.2018.00248

Besedovsky, L., Ngo, H.-V. V., Dimitrov, S., Gassenmaier, C., Lehmann, R., & Born, J. (2017). Auditory closed-loop stimulation of EEG slow oscillations strengthens sleep and signs of its immune-supportive function. Nature Communications, 8(1), 1984. https://doi.org/10.1038/s41467-017-02170-3

Bigdely-Shamlo, N., Mullen, T., Kothe, C., Su, K.-M., & Robbins, K. A. (2015). The PREP pipeline: standardized preprocessing for large-scale EEG analysis. Frontiers in Neuroinformatics, 0(JUNE), 16. https://doi.org/10.3389/FNINF.2015.00016

Brandenberger, G., Ehrhart, J., Piquard, F., & Simon, C. (2001). Inverse coupling between ultradian oscillations in delta wave activity and heart rate variability during sleep. Clinical Neurophysiology, 112(6), 992–996. https://doi.org/10.1016/S1388-2457(01)00507-7

Cajochen, C., Pischke, J., Aeschbach, D., & Borbély, A. A. (1994). Heart rate dynamics during human sleep. Physiology and Behavior, 55(4), 769–774. https://doi.org/10.1016/0031-9384(94)90058-2

Carter, M. E., Yizhar, O., Chikahisa, S., Nguyen, H., Adamantidis, A., Nishino, S., Deisseroth, K., & De Lecea, L. (2010). Tuning arousal with optogenetic modulation of locus coeruleus neurons. Nature Neuroscience 2010 13:12, 13(12), 1526–1533. https://doi.org/10.1038/nn.2682

Chaieb, L., Wilpert, E. C., Reber, T. P., & Fell, J. (2015). Auditory beat stimulation and its effects on cognition and mood states. In Frontiers in Psychiatry (Vol. 6, Issue MAY). Frontiers Media S.A. https://doi.org/10.3389/fpsyt.2015.00070

Delorme, A., & Makeig, S. (2004). EEGLAB: an open source toolbox for analysis of single-trial EEG dynamics including independent component analysis. J Neurosci Methods, 134(1), 9–21.

Diekelmann, S., & Born, J. (2010). The memory function of sleep. Nature Reviews Neuroscience, 11(2), 114–126. https://doi.org/10.1038/nrn2762

Dijk, D.-J. J. (2009). Regulation and functional correlates of slow wave sleep. Journal of Clinical Sleep Medicine: JCSM: Official Publication of the American Academy of Sleep Medicine, 5(2 Suppl), S6. https://doi.org/10.5664/jcsm.5.2s.s6

Dijk, D. J. (2008). Slow-wave sleep, diabetes, and the sympathetic nervous system. Proceedings of the National Academy of Sciences of the United States of America. https://doi.org/10.1073/pnas.0711635105

Eide, P. K., Vinje, V., Pripp, A. H., Mardal, K.-A., & Ringstad, G. (2021). Sleep deprivation impairs molecular clearance from the human brain. Brain, 144(3), 863–874. https://doi.org/10.1093/brain/awaa443

Esser, S. K., Hill, S. L., & Tononi, G. (2007). Sleep homeostasis and cortical synchronization: I. Modeling the effects of synaptic strength on sleep slow waves. Sleep, 30(12), 1617–1630. https://doi.org/10.1093/sleep/30.12.1617

Eugene, A. R., & Masiak, J. (2015). The Neuroprotective Aspects of Sleep. MEDtube Science.

Fattinger, S., De Beukelaar, T. T., Ruddy, K. L., Volk, C., Heyse, N. C., Herbst, J. A., Hahnloser, R. H. R., Wenderoth, N., & Huber, R. (2017). Deep sleep maintains learning efficiency of the human brain. Nature Communications, 8(May), 1–13. https://doi.org/10.1038/ncomms15405

Fernández-Varela, I., Alvarez-Estevez, D., Hernández-Pereira, E., & Moret-Bonillo, V. (2017). A simple and robust method for the automatic scoring of EEG arousals in polysomnographic recordings. Computers in Biology and Medicine, 87, 77–86.

Ferrarelli, F., Huber, R., Peterson, M. J., Massimini, M., Murphy, M., Riedner, B. A., Watson, A., Bria, P., & Tononi, G. (2007). Reduced sleep spindle activity in schizophrenia patients. The American Journal of Psychiatry, 164(3), 483–492. https://doi.org/10.1176/AJP.2007.164.3.483

Ferrarelli, F., Peterson, M. J., Sarasso, S., Riedner, B. A., Murphy, M. J., Benca, R. M., Bria, P., Kalin, N. H., & Tononi, G. (2010). Thalamic dysfunction in schizophrenia suggested by whole-night deficits in slow and fast spindles. The American Journal of Psychiatry, 167(11), 1339–1348. https://doi.org/10.1176/APPI.AJP.2010.09121731

Ferster, M. L., Lustenberger, C., & Karlen, W. (2019). Configurable mobile system for autonomous high-quality sleep monitoring and closed-loop acoustic stimulation. IEEE Sensors Letters, 3(5), 1–1. https://doi.org/10.1109/LSENS.2019.2914425

Foote, S. L., & Morrison, J. H. (1987). Extrathalamic modulation of cortical function. Annual Review of Neuroscience, 10, 67–95. https://doi.org/10.1146/ANNUREV.NE.10.030187.000435

Grimaldi, D., Papalambros, N. A., Reid, K. J., Abbott, S. M., Malkani, R. G., Gendy, M., Iwanaszko, M., Braun, R. I., Sanchez, D. J., Paller, K. A., & Zee, P. C. (2019). Strengthening sleep-autonomic interaction via acoustic enhancement of slow oscillations. Sleep, February, 1–11. https://doi.org/10.1093/sleep/zsz036

Helfrich, R. F., Mander, B. A., Jagust, W. J., Knight, R. T., & Walker, M. P. (2018). Old Brains Come Uncoupled in Sleep: Slow Wave-Spindle Synchrony, Brain Atrophy, and Forgetting. Neuron, 97(1), 221-230.e4. https://doi.org/10.1016/j.neuron.2017.11.020

Iber, C., Ancoli-Israel, S., Chesson, A., & others. (2007). The AASM Manual for the Scoring of Sleep and Associated Events - Rules Terminlogy and Technical Specifications. American A.

Jurysta, F., Van De Borne, P., Migeotte, P. F., Dumont, M., Lanquart, J. P., Degaute, J. P., & Linkowski, P. (2003). A study of the dynamic interactions between sleep EEG and heart rate variability in healthy young men. Clinical Neurophysiology, 114(11), 2146–2155. https://doi.org/10.1016/S1388-2457(03)00215-3

Kearney, K. (n.d.). boundedline.m (https://github.com/kakearney/boundedline-pkg). GitHub. Retrieved December 9, 2021, from https://github.com/kakearney/boundedline-pkg

Knutson, K. L., Spiegel, K., Penev, P., & Van Cauter, E. (2007). The metabolic consequences of sleep deprivation. Sleep Med Rev, 11(3), 163–178. https://doi.org/10.1016/j.smrv.2007.01.002

Kraus, N., McGee, T., Littman, T., Nicol, T., & King, C. (1994). Nonprimary auditory thalamic representation of acoustic change. https://Doi.Org/10.1152/Jn.1994.72.3.1270, 72(3), 1270–1277. https://doi.org/10.1152/JN.1994.72.3.1270

Kuznetsova, A., Brockhoff, P. B., Christensen, R. H. B., & Jensen, S. P. (2020). Package ‘ lmerTest .’ https://github.com/runehaubo/lmerTestR

Lakatta, E. G. (2003). Arterial and Cardiac Aging: Major Shareholders in Cardiovascular Disease Enterprises. Circulation, 107(3), 490–497. https://doi.org/10.1161/01.cir.0000048894.99865.02

Landolt, H. P., Dijk, D. J., Achermann, P., & Borbély, A. A. (1996). Effect of age on the sleep EEG: Slow-wave activity and spindle frequency activity in young and middle-aged men. Brain Research, 738(2), 205–212. https://doi.org/10.1016/S0006-8993(96)00770-6

Landsness, E. C., Crupi, D., Hulse, B. K., Peterson, M. J., Huber, R., Ansari, H., Coen, M., Cirelli, C., Benca, R. M., Ghilardi, M. F., & Tononi, G. (2009). Sleep-dependent improvement in visuomotor learning: A causal role for slow waves. Sleep, 32(10), 1273–1284. https://doi.org/10.1093/sleep/32.10.1273

Leminen, M. M., Virkkala, J., Saure, E., Paajanen, T., Zee, P. C., & Santostasi, G. (2017). Enhanced Memory Consolidation Via Automatic Sound Stimulation During Non-REM Sleep. 40(3). https://doi.org/10.1093/sleep/zsx003

Lustenberger, C., Patel, Y. A., Alagapan, S., Page, J. M., Price, B., Boyle, M. R., & Fröhlich, F. (2018). High-density EEG characterization of brain responses to auditory rhythmic stimuli during wakefulness and NREM sleep. NeuroImage, 169(August 2017), 57–68. https://doi.org/10.1016/j.neuroimage.2017.12.007

Lustenberger, C., Wehrle, F., Tüshaus, L., Achermann, P., & Huber, R. (2015). The Multidimensional Aspects of Sleep Spindles and Their Relationship to Word-Pair Memory Consolidation. Sleep, 38(7), 1093–1103. https://doi.org/10.5665/sleep.4820

Massimini, M., Huber, R., Ferrarelli, F., Hill, S., & Tononi, G. (2004). The sleep slow oscillation as a traveling wave. The Journal of Neuroscience : The Official Journal of the Society for Neuroscience, 24(31), 6862–6870. https://doi.org/10.1523/JNEUROSCI.1318-04.2004

McBurney-Lin, J., Lu, J., Zuo, Y., & Yang, H. (2019). Locus coeruleus-norepinephrine modulation of sensory processing and perception: A focused review. Neuroscience & Biobehavioral Reviews, 105(3), 190–199. https://doi.org/10.1016/j.neubiorev.2019.06.009

Mullington, J. M., Haack, M., Toth, M., Serrador, J. M., & Meier-Ewert, H. K. (2009). Cardiovascular, inflammatory, and metabolic consequences of sleep deprivation. Progress in Cardiovascular Diseases, 51(4), 294–302.

Ngo Hong-Viet V., Claussen, J. C., Born, J., & Mölle, M. (2013). Induction of slow oscillations by rhythmic acoustic stimulation. Journal of Sleep Research, 22(1), 22–31. https://doi.org/10.1111/j.1365-2869.2012.01039.x

Ngo Hong-Viet V., Martinetz, T., Born, J., & Mölle, M. (2013). Auditory Closed-Loop Stimulation of the Sleep Slow Oscillation Enhances Memory. Neuron, 78(3), 545–553. https://doi.org/10.1016/j.neuron.2013.03.006

Ngo, H.-V. V, Miedema, A., Faude, I., Martinetz, T., Molle, M., & Born, J. (2015). Driving Sleep Slow Oscillations by Auditory Closed-Loop Stimulation--A Self-Limiting Process. Journal of Neuroscience, 35(17), 6630–6638. https://doi.org/10.1523/JNEUROSCI.3133-14.2015

Nunan, D., Sandercock, G. R. H., & Brodie, D. A. (2010). A Quantitative Systematic Review of Normal Values for Short-Term Heart Rate Variability in Healthy Adults. Pacing and Clinical Electrophysiology, 33(11), 1407–1417. https://doi.org/10.1111/j.1540-8159.2010.02841.x

Ong, J. L., Lo, J. C., Chee, N. I. Y. N., Santostasi, G., Paller, K. A., Zee, P. C., & Chee, M. W. L. (2016). Effects of phase-locked acoustic stimulation during a nap on EEG spectra and declarative memory consolidation. Sleep Medicine, 20, 88–97.

Ong, J. L., Patanaik, A., Chee, N. I. Y. N., Lee, X. K., Poh, J.-H., & Chee, M. W. L. (2018). Auditory stimulation of sleep slow oscillations modulates subsequent memory encoding through altered hippocampal function. Sleep, 41(5), zsy031.

Otzenberger, H., Gronfier, C., Simon, C., Charloux, A., Ehrhart, J., Piquard, F., & Brandenberger, G. (1998). Dynamic heart rate variability: A tool for exploring sympathovagal balance continuously during sleep in men. American Journal of Physiology - Heart and Circulatory Physiology, 275(3 44-3). https://doi.org/10.1152/ajpheart.1998.275.3.h946

Palagini, L., Bruno, R. M., Gemignani, A., Baglioni, C., Ghiadoni, L., & Riemann, D. (2013). Sleep loss and hypertension: a systematic review. Curr Pharm Des, 19(13), 2409–2419. http://www.ncbi.nlm.nih.gov/pubmed/23173590

Papalambros, N. A., Santostasi, G., Malkani, R. G., Braun, R., Weintraub, S., Paller, K. A., & Zee, P. C. (2017). Acoutstic enhancement of sleep slow oscillations and concomitant memory improvement in older adults. Frontiers in Human Neuroscience, 11(March), 1–14. https://doi.org/10.3389/fnhum.2017.00109

Papalambros, N. A., Weintraub, S., Chen, T., Grimaldi, D., Santostasi, G., Paller, K. A., Zee, P. C., & Malkani, R. G. (2019). Acoustic enhancement of sleep slow oscillations in mild cognitive impairment. Annals of Clinical and Translational Neurology, 6(7), 1191–1201.

Parrino, L., & Vaudano, A. E. (2018). The resilient brain and the guardians of sleep: New perspectives on old assumptions. Sleep Medicine Reviews, 39, 98–107. https://doi.org/10.1016/j.smrv.2017.08.003

Pivik, R. T., & Harman, K. (1995). A reconceptualization of EEG alpha activity as an index of arousal during sleep: all alpha activity is not equal. Journal of Sleep Research, 4(3), 131–137. https://doi.org/10.1111/j.1365-2869.1995.tb00161.x

R Core Team. (2013). R: A language and environment for statistical computing.

Renard, Y., Lotte, F., Gibert, G., Congedo, M., Maby, E., Delannoy, V., Bertrand, O., & Lécuyer, A. (2010). OpenViBE: An open-source software platform to design, test, and use brain-computer interfaces in real and virtual environments. Presence: Teleoperators and Virtual Environments, 19(1), 35–53. https://doi.org/10.1162/pres.19.1.35

Riedner, B. A., Vyazovskiy, V. V., Huber, R., Massimini, M., Esser, S., Murphy, M., & Tononi, G. (2007). Sleep homeostasis and cortical synchronization: III. A high-density EEG study of sleep slow waves in humans. Sleep, 30(12), 1643–1657. https://doi.org/10.1093/sleep/30.12.1643

Santostasi, G., Malkani, R., Riedner, B., Bellesi, M., Tononi, G., Paller, K. A., & Zee, P. C. (2015). Phase-locked loop for precisely timed acoustic stimulation during sleep. Journal of Neuroscience Methods, 259, 101–114. https://doi.org/10.1016/j.jneumeth.2015.11.007

Schlosser, J. (1996). Heart rate variability: standards of measurement, physiological interpretation and clinical use. Task Force of the European Society of Cardiology and the North American Society of Pacing and Electrophysiology. Circulation, 93(5), 1043–1065. https://doi.org/10.1161/01.CIR.93.5.1043

Searle, S. R., Speed, F. M., & Milliken, G. A. (2021). Estimated Marginal Means, aka Least-Squares Means [R package emmeans version 1.7.1-1]. American Statistician, 34(4), 216–221. https://doi.org/10.1080/00031305.1980.10483031

Shaffer, F., & Ginsberg, J. P. (2017). An Overview of Heart Rate Variability Metrics and Norms. Frontiers in Public Health, 5(September), 1–17. https://doi.org/10.3389/fpubh.2017.00258

Simon, C., Gronfier, C., Schlienger, J. L., & Brandenberger, G. (1998). Circadian and ultradian variations of leptin in normal man under continuous enteral nutrition: Relationship to sleep and body temperature. Journal of Clinical Endocrinology and Metabolism. https://doi.org/10.1210/jcem.83.6.4864

Simor, P., Steinbach, E., Nagy, T., Gilson, M., Farthouat, J., Schmitz, R., Gombos, F., Ujma, P. P., Pamula, M., Bódizs, R., & Peigneux, P. (2018). Lateralized rhythmic acoustic stimulation during daytime NREM sleep enhances slow waves. Sleep, 41(12). https://doi.org/10.1093/sleep/zsy176

Somers, V. K., Dyken, M. E., Mark, A. L., & Abboud, F. M. (1993). Sympathetic-Nerve Activity during Sleep in Normal Subjects. New England Journal of Medicine. https://doi.org/10.1056/NEJM199302043280502

Steriade, M. (2006). Grouping of brain rhythms in corticothalamic systems. Neuroscience, 137(4), 1087–1106. https://doi.org/10.1016/J.NEUROSCIENCE.2005.10.029

Steriade, M., Nuñez, A., & Amzica, F. (1993). A novel slow inferior to 1 Hz oscillation of neocortical neurons in vivo: depolarizing and hyperpolarizing components. The Journal of Neuroscience : The Official Journal of the Society for Neuroscience.

Stich, F. M., Huwiler, S., D’Hulst, G., & Lustenberger, C. (2021). The potential role of sleep in promoting a healthy body composition: Underlying mechanisms determining muscle, fat, and bone mass and their association to sleep. Neuroendocrinology. https://doi.org/10.1159/000518691

Tasali, E., Leproult, R., Ehrmann, D. A., & Van Cauter, E. (2008). Slow-wave sleep and the risk of type 2 diabetes in humans. Proceedings of the National Academy of Sciences of the United States of America. https://doi.org/10.1073/pnas.0706446105

Thayer, J. F., Yamamoto, S. S., & Brosschot, J. F. (2010). The relationship of autonomic imbalance, heart rate variability and cardiovascular disease risk factors. International Journal of Cardiology, 141(2), 122–131. https://doi.org/10.1016/j.ijcard.2009.09.543

Thong, T., Li, K., McNames, J., Aboy, M., & Goldstein, B. (2003). Accuracy of Ultra-Short Heart Rate Variability Measures. Annual International Conference of the IEEE Engineering in Medicine and Biology - Proceedings, 3(March 2014), 2424–2427. https://doi.org/10.1109/IEMBS.2003.1280405

Tononi, G., & Cirelli, C. (2006). Sleep function and synaptic homeostasis. In Sleep Medicine Reviews. https://doi.org/10.1016/j.smrv.2005.05.002

Underwood, E. (2013). Sleep: The brain’s housekeeper? In Science. https://doi.org/10.1126/science.342.6156.301

Weigenand, A., Mölle, M., Werner, F., Martinetz, T., & Marshall, L. (2016). Timing matters: open-loop stimulation does not improve overnight consolidation of word pairs in humans. European Journal of Neuroscience, 44(6), 2357–2368. https://doi.org/10.1111/ejn.13334

WHO. (2016). Cardiovascular diseases (CVDs) fact sheets. Who.

Wickham, H. (2016). ggplot2: Elegant Graphics for Data Analysis. Springer-Verlag New York. https://ggplot2.tidyverse.org

Wunderlin, M., Züst, M. A., Hertenstein, E., Fehér, K. D., Schneider, C. L., Klöppel, S., & Nissen, C. (2021). Modulating overnight memory consolidation by acoustic stimulation during slow wave sleep – a systematic review and meta-analysis. Sleep, 46(May), 1–17. https://doi.org/10.1093/sleep/zsaa296

Xi, B., He, D., Zhang, M., Xue, J., & Zhou, D. (2014). Short sleep duration predicts risk of metabolic syndrome: a systematic review and meta-analysis. Sleep Medicine Reviews, 18(4), 293–297.

