## Appendix for "Effects of auditory sleep modulation approaches on brain oscillatory and cardiovascular dynamics"

### Table A1: Overview of different stimulation conditions

| Study | Stimulation Modality | Category | Description |
|---|---|---|---|
| 1 | UP | Phase-specific | 50 ms pink noise targeting ascending phase of slow waves |
| 1 | DOWN | Phase-specific | 50 ms pink noise targeting descending phase of slow waves |
| 1,2,3 | $ISI1_{High}$ | Rhythmic | 1 Hz rhythmic stimulation at 45 dB |
| 1 | ENVELOPE | Sound modulation | 1 Hz amplitude modulated continuous pink noise |
| 1 | BINAURAL BEATS | Sound modulation | Binaural beats played with 1 Hz frequency difference and carrier frequency of 400 Hz |
| 2 | $ISI1_{Mod}$ | Rhythmic | 1 Hz rhythmic stimulation with increasing and decreasing sound volume |
| 3 | $ISI1_{Low}$ | Rhythmic | 1 Hz rhythmic stimulation at 42.5 dB |
| 1,2,3 | SHAM | - | Control condition: no tones are played |

### Table A2: Sleep and stimulation number descriptive statistics for studies 1,2, and 3

Table shows sleep architecture and basic stimulation statistics (number stimulation windows occurring within the first four hours after stimulation started: early night, and number stimulation windows occurring in the remaining hours of the night: late night). Total sleep time (TST), wake after sleep onset (WASO), non-rapid eye movement (N) sleep stages 1 (N1), 2 (N2) and 3 (N3), rapid-eye movement sleep (REM). Latency refers to the first occurrence of the mentioned sleep stage. All data is presented as mean ± standard error of the mean.

| | **Study1 (n = 23)** | **Study2 (n = 9)** | **Study3 (n = 19)** |
|---|---|---|---|
| **Number stimulation windows early night** | 199.13 ± 23.46 | 307.22 ± 34.22 | 300.16 ± 22.35 |
| **Number stimulation windows late night** | 118.13 ± 17.65 | 103.22 ± 26.08 | 155.47 ± 19.75 |
| **First stimulation [min]** | 44.35 ± 6.21 | 60.17 ± 23.26 | 49.26 ± 7.44 |
| **Sleep efficiency [a.u.]** | 85.63 ± 1.67 | 87.35 ± 2.65 | 87.13 ± 1.24 |
| **TST [min]** | 349.65 ± 5.51 | 358.19 ± 8.58 | 345.21 ± 9.87 |
| **WASO [min]** | 60.07 ± 8.24 | 51.74 ± 11.74 | 52.245 ± 7.62 |
| **Wake [min]** | 69.97 ± 8.19 | 61.22 ± 12.95 | 61.96 ± 7.65 |
| **N1 [min]** | 81.65 ± 7.46 | 66.07 ± 7.54 | 53.12 ± 5.42 |
| **N2 [min]** | 224.49 ± 4.10 | 248.48 ± 20.15 | 239.84 ± 11.50 |
| **N3 [min]** | 43.17 ± 6.27 | 43.30 ± 12.81 | 51.87 ± 7.24 |
| **REM [min]** | 69.84 ± 5.43 | 66.67 ± 7.41 | 77.54 ± 7.18 |
| **N2 latency [min]** | 19.84 ± 3.27 | 14.30 ± 2.17 | 19.84 ± 2.26 |
| **REM latency [min]** | 147.81 ± 17.31 | 148.63 ± 11.67 | 116.07 ± 15.44 |

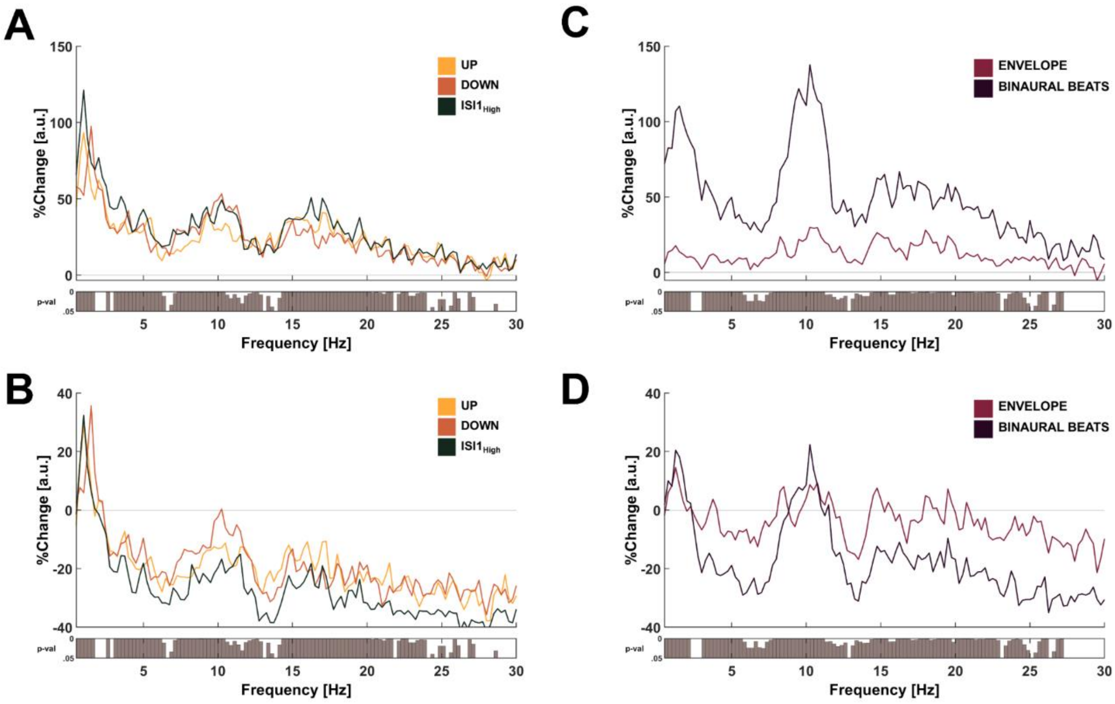


**Figure A1: Relative power spectral density distribution during ON window**.
A + C: Percentage change of power spectral density distribution for conditions UP, DOWN, ($ISI1_{High}$) (A) or ENVELOPE and BINAURAL BEATS (B) to SHAM condition. Note that for these plots the non-normalized power spectrum in bins with a resolution of 0.25 was used for comparisons. The horizontal line at 0 represents no change compared to the SHAM condition. Below the spectrum, the resulting p-values of a linear mixed-effects model for each bin with the fixed factor condition and random factor participants are shown.
B + D: Percentage change of normalized power spectral density distribution for conditions UP, DOWN, $ISI1_{High}$ (C) or ENVELOPE and BINAURAL BEATS (D) to SHAM condition. Note that for these we normalized the power spectrum from 0.5 to 30 Hz and the bin resolution of 0.25 was used for comparisons. The horizontal line at 0 represents no change compared to the SHAM condition. Below the spectrum, the resulting p-values of a linear mixed-effects model for each bin with the fixed factor condition and random factor participants are shown. Data for n = 23 participants is shown.

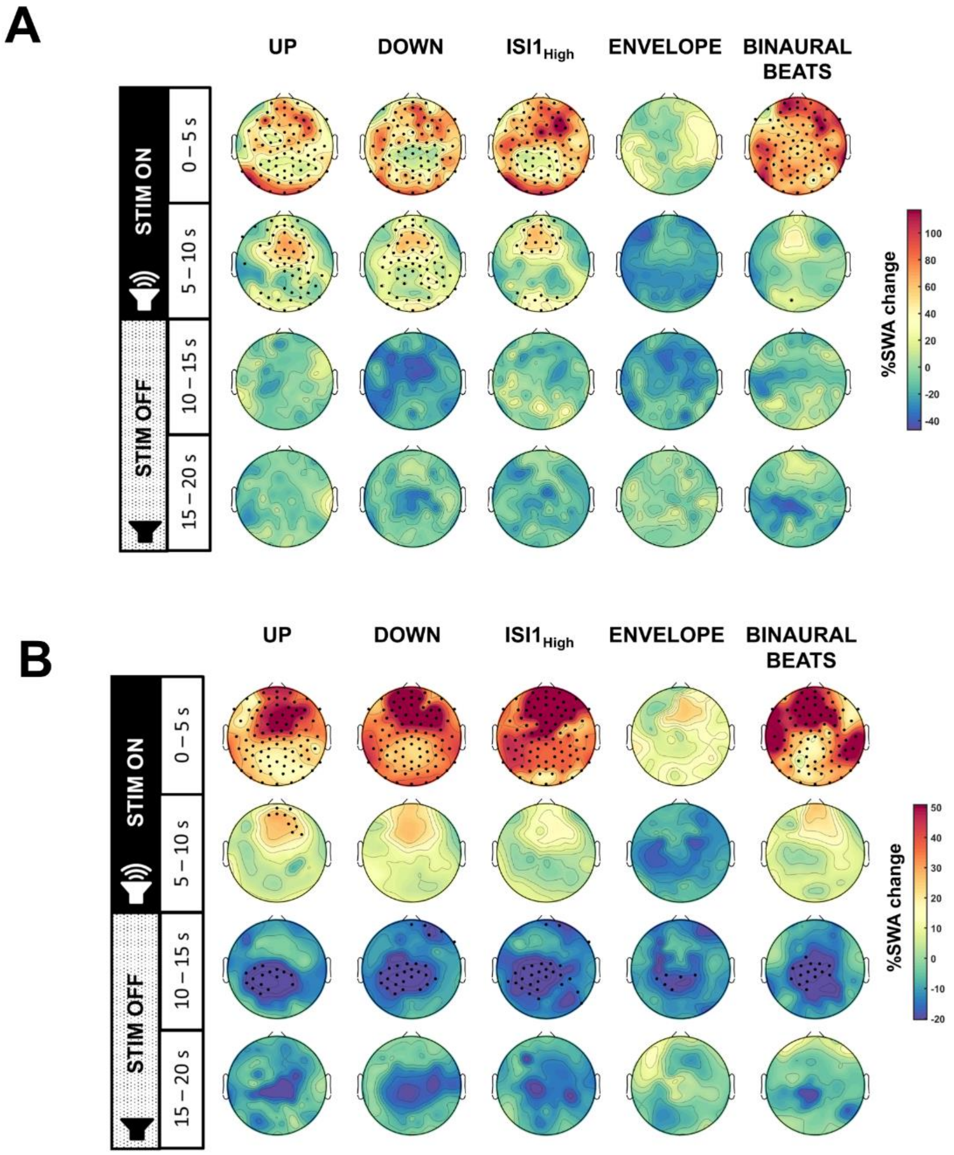


**Figure A2: Effects of auditory stimulation conditions on change in slow wave activity compared to SHAM.**
A + B: Percentage change of slow wave activity (SWA, A: 0.5 – 2 Hz, B: high SWA: 2.25 - 4.5 Hz) for the conditions UP, DOWN, $ISI1_{High}$, ENVELOPE and BINAURAL BEATS compared to SHAM. We divided the stimulation ON and OFF window in two 5 s windows each. Black dots indicate significant electrodes ($p < 0.05$) for the post-hoc p-values resulting from linear mixed-effects effects model models with condition as fixed factor and subject as random factor. P-values for each topoplot have been corrected for multiple comparisons by applying the false discovery rate. All plots show data for for n = 23 participants.

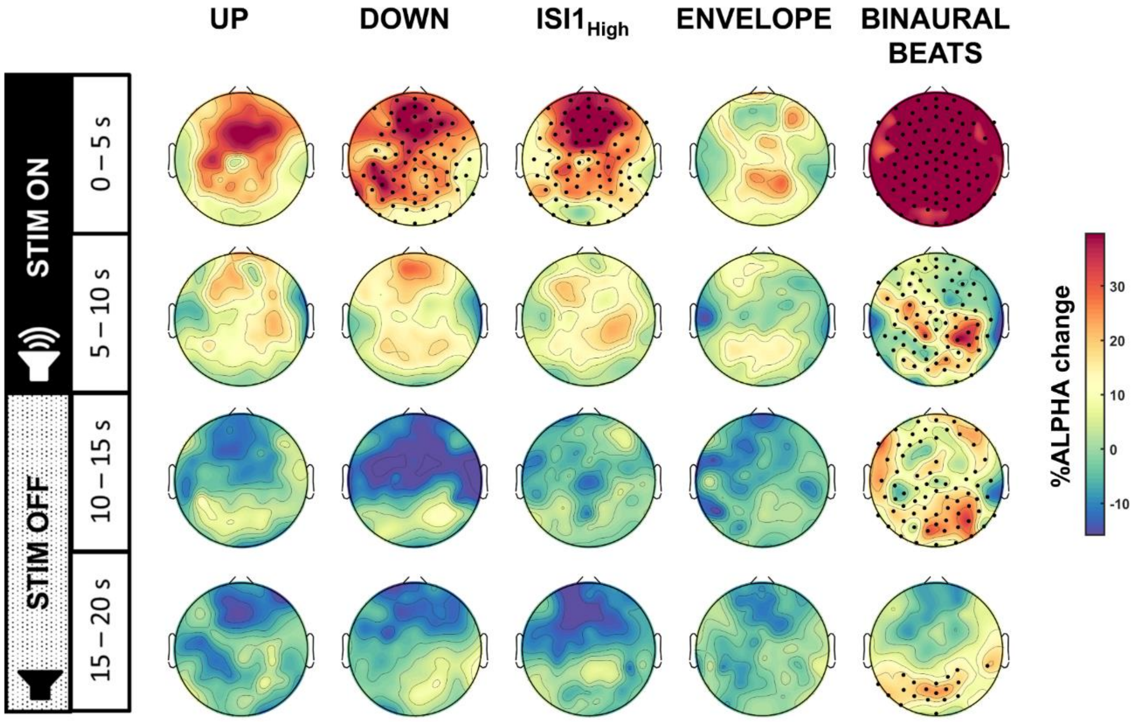


**Figure A3: Effects of auditory stimulation conditions on change in low alpha activity compared to SHAM.**
Percentage change of low alpha activity (8 - 10 Hz) for the conditions UP, DOWN, $ISI1_{High}$, ENVELOPE and BINAURAL BEATS compared to SHAM. We divided the stimulation ON and OFF window in two 5 s windows each. Black dots indicate significant electrodes ($p < 0.05$) for the post-hoc p-values resulting from linear mixed-effects effects model models with condition as fixed factor and subject as random factor. P-values for each topoplot have been corrected for multiple comparisons by applying the false discovery rate. All plots are shown for n = 23 participants

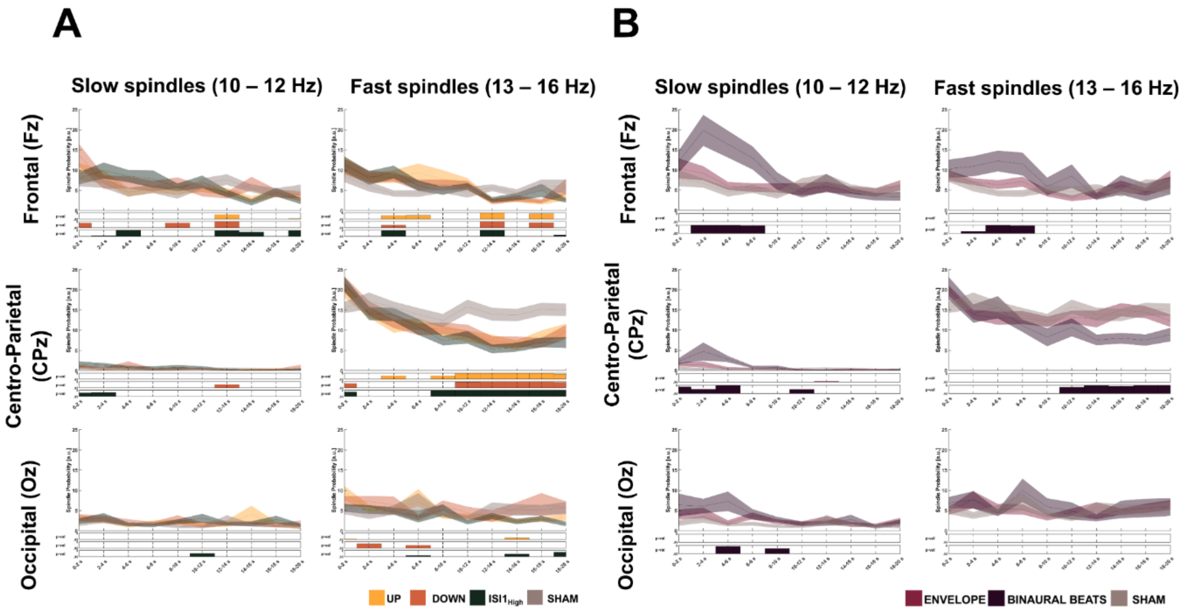


**Figure A4: Probability of spindles to occur during stimulation windows.**
A + B: Probability of an occurring spindle start in 2 seconds bins of the stimulation window. Probability data is presented as mean ± standard error of the mean. Resulting p-values of linear mixed-effects models within each bin for condition as fixed factor and subject as random factor are shown below the graph in yellow (UP), orange (DOWN), dark green ($ISI1_{High}$), and grey (SHAM) for A, or pink (ENVELOPE) and dark violet (BINAURAL BEATS), and grey (SHAM) in B, respectively. We divided spindles in slow spindles (10 - 12 Hz) and fast spindles (13 - 16 Hz) and plotted the probability for the Fz, CPz, and Oz electrode for n = 23 participants.

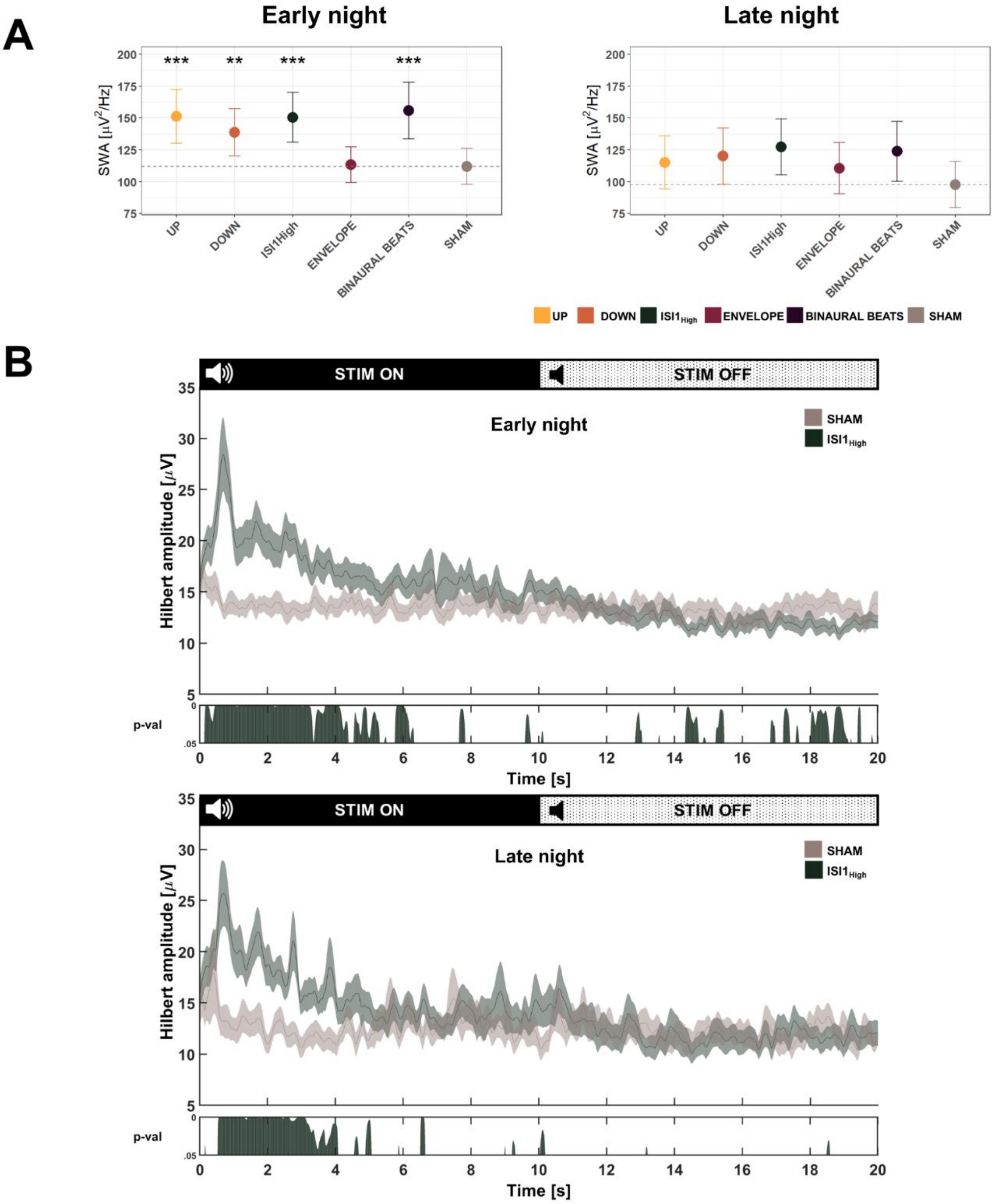


**Figure A3: Comparison of brain response to auditory stimulation during the first four hours after stimulation started (early night) and the remaining hours of the stimulation (late night).**

A: Slow wave activity (SWA, 0.5 – 2 Hz) for the stimulation window (ONOFF together) for the conditions UP, DOWN, $ISI1_{High}$, ENVELOPE, BINAURAL BEATS, and SHAM for n = 23 participants. Post-hoc comparisons of a linear mixed-effects model entering condition as fixed factor and subject as random factor were corrected for multiple comparisons using the Hochberg method. ***: $p < 0.001$, **: $p < 0.01$, *: $p < 0.05$, #: $0.05 < p < 0.1$.

B: Hilbert amplitude response (0.5 – 2 Hz) for the $ISI1_{High}$ and SHAM stimulation during the stimulation window for the early night (top) and the late night (bottom). Below the plot the resulting p-values of the post-hoc comparison of linear mixed effects-models with condition as fixed factor and subject as random factor for each time point are shown. Data is presented as mean ± standard error of the mean for n = 23 participants.

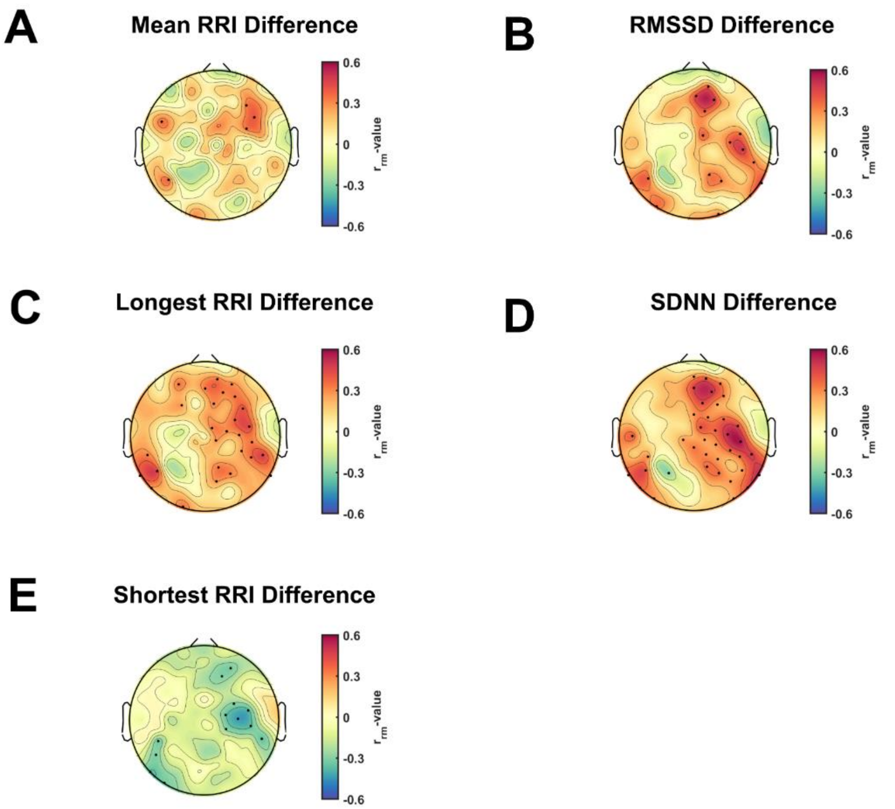


**Figure A4: Repeated measures correlations** between percentage change for heart rate (variability; HR(V)) features and percentage change of alpha activity (8 - 10 Hz) in the first five seconds of the stimulation ON window. For every participant (n = 22) and every condition a mean value for each variable was calculated. We excluded one participant because not all conditions had ECG measurements of high enough qualities. Topoplots show r values and significant electrodes ($p < 0.05$) are marked with black dots. We corrected the p-values for multiple comparisons using false discovery rate. A: Mean difference of consecutive heart beats (RRI). B: Difference in the root mean square of successive difference of normal heart beats (RMSSD). C: Differences of the longest RRI, D: Difference in the standard deviation of differences between normal heartbeats (SDNN). E: Percentage of difference of the shortest RRI correlated with percentage change in SWA for electrode Pz.
